# Plastin-3 membrane recruitment drives cell-in-cell invasion during entosis

**DOI:** 10.64898/2026.03.17.709257

**Authors:** Stavroula Prapa, Emir Bozkurt, Carolina Gandia, Alberto Hernandez-Cano, Mario Soriano, Luke A. Noon, Jose A Gomez-Sanchez, Mar Orzáez Calatayud, Federico Lucantoni

**Affiliations:** Cellular Stress and Cell Death Pathways Laboratory, Centro de Investigación Príncipe Felipe (CIPF), Valencia, Spain; Targeted Therapies on Cancer and Inflammation Laboratory, Centro de Investigación Príncipe Felipe (CIPF), Valencia, Spain; Department of Mechanical Engineering, Virginia Tech, Blacksburg, U.S.A; Servicio de Microscopía Óptica Avanzada, Centro de Investigación Príncipe Felipe (CIPF), Valencia, Spain; Servicio de Microscopía Electrónica, Centro de Investigación Príncipe Felipe (CIPF), Valencia, Spain; Metabolic Growth Signals and Regenerative Medicine Laboratory, Centro de Investigación Príncipe Felipe (CIPF), Valencia, Spain; Instituto de Investigación Sanitaria y Biomédica de Alicante (ISABIAL), Alicante, Spain; Unidad Mixta UPV-CIPF de Investigación en Mecanismos de Enfermedades y Nanomedicina, Universitat Politècnica de València, Centro de Investigación Príncipe Felipe, Valencia, Spain

**Keywords:** Entosis, Cell-in-Cell, ROCK1, Plastin-3, Breast cancer

## Abstract

Entosis is a cell-in-cell (CIC) invasion process in which one living cell actively invades into another, with important implications for tumor evolution, cell competition, and responses to cellular stress. Both the initiation and progression of entosis require extensive actin cytoskeletal remodeling, as one cell must physically accommodate the complete internalization of another viable cell, which can subsequently migrate, divide, undergo degradation, or escape. Nearly two decades after its discovery, mechanistic insight into how cytoskeleton organization is regulated during entosis remains poorly understood. The most consistent observation is that inhibition of Rho–ROCK signaling blocks entosis, indicating a central role for actomyosin contractility. Consistently, increased phosphorylation of myosin light chain and cortical actomyosin accumulation have been reported in invading cells, yet the underlying mechanisms remain largely unknown.

Here, we systematically interrogated ROCK activation as a driver of the entotic internalization program. Constitutively active mutant ROCK1 or pharmacological ROCK1 activation using narciclasine rapidly triggered canonical entosis pathway through MLC2 phosphorylation. Mass spectrometry identified Plastin-3 (PLS3, T-plastin), a crucial calcium-sensitive actin-bundling protein, which was recruited to the plasma membrane to trigger entosis upon ROCK1 activation. Our study uncovers a key cytoskeletal remodeling step which may support cortical tension required for CIC invasion during entosis.

## INTRODUCTION

Entosis is a conserved cell invasion process in which a living cell actively invades into a neighboring cell, generating characteristic cell-in-cell (CIC) structures^1^ with diverse biological consequences^2,3^. This phenomenon has emerged as an important determinant of tumor evolution, cell competition, and therapeutic response, influencing clonal selection, genome stability, immune evasion and patient prognosis in cancer^4–11^.

The initiation of entosis relies on cytoskeletal remodeling controlled by the RhoA–ROCK1 signaling axis. RhoA, a member of the small GTPase family, activates the serine/threonine kinase ROCK1, which regulates actomyosin contractility through phosphorylation of downstream effectors such as myosin light chain (MLC)^12,13^. Despite these observations, the molecular factors that couple ROCK-dependent contractility to the physical process of membrane remodeling and cell internalization are still largely unknown, leaving the signaling events that trigger the initiation of entosis unresolved. The reason of this lack of information is that entosis typically occurs at low frequency, which has limited detailed molecular dissection of the process. Moreover, it is most commonly observed under indirect stress conditions^14^, including glucose deprivation^15^, loss of matrix attachment^16^, aberrant mitosis^17^ or exposure to cytotoxic stimuli^18,19^. These contexts limit precise mechanistic interrogation and raise the unresolved question of whether entosis represents a directly modulable cellular behavior or instead arise secondarily from cellular stress or damage.

Establishing direct activation of entosis is therefore critical for defining its mechanistic logic and biological function. In particular, although ROCK1 activity is clearly required for entotic internalization, whether activation of the ROCK1 pathway alone is sufficient to initiate entosis remains unclear. Additionally, during entosis, the host cell must reorganize large portions of its cytoskeleton to accommodate a cell of comparable size, requiring extensive remodeling of the actin network and the coordinated activity of multiple cytoskeletal regulators, which are unknown.

Resolving this question would 1) distinguish ROCK1 as a permissive factor from a true triggering signal, 2) provide an experimental entry point to manipulate entosis independently of confounding stress responses, and 3) identify additional proteins and signaling pathways that support actin assembly, contractility, and membrane remodeling.

Here, we identify constitutive active mutant ROCK1 or pharmacological activation of ROCK1 as sufficient to rapidly trigger entosis through induction of actomyosin contractility. Acute ROCK1 activation increases myosin light-chain phosphorylation and promotes entotic structure formation within 6 hours, whereas pharmacological inhibition or genetic silencing of ROCK1 abolishes this response.

Mechanistically, we further identify Plastin-3 (PLS3), a calcium dependent actin-bundling protein implicated in cytoskeletal organization and cancer progression, as an essential regulator of ROCK1-driven entosis; upon ROCK1 activation, PLS3 accumulated at the plasma membrane. Functionally, PLS3 overexpression increased entotic formation, whereas PLS3 silencing markedly reduced the entosis induced by narciclasine. This work provides a mechanistic framework and tractable platform for interrogating the roles of entosis in tumor biology and therapeutic response.

## MATERIALS AND METHODS

### Materials and reagents

Fetal bovine serum (FBS), RPMI-1640 medium, and dimethyl sulfoxide (DMSO) were purchased from Sigma-Aldrich (St. Louis, MO, USA). Narciclasine, Y27632, zVAD-fmk, and Blebbistatin were obtained from MedChemExpress (Monmouth Junction, NJ, USA). The SiR-actin kit was purchased from Spirochrome (Stein am Rhein, Switzerland). Hoechst 33342, ProLong™ Diamond Antifade Mountant with DAPI, LysoTracker™ Red DND-99, LysoTracker™ Deep Red, Opti-MEM™ Reduced Serum Medium, FluoroBrite™ DMEM and Lipofectamine™ 2000 were obtained from Thermo Fisher Scientific (Waltham, MA, USA). Glass-bottom WillCo dishes were purchased from WillCo Wells B.V. (Amsterdam, The Netherlands).

### Cell lines

MCF7, EFM19, ZR75-1, MCF10A, and HeLa cell lines were maintained under standard culture conditions at 37 °C in a humidified incubator with 5% CO₂. MCF7 and EFM19 cells were cultured in RPMI-1640 medium supplemented with 10% fetal bovine serum (FBS) and 1% penicillin–streptomycin. ZR75-1 cells were maintained in DMEM/F-12 medium supplemented with 10% FBS and 1% penicillin–streptomycin. HeLa cells were cultured in high-glucose DMEM supplemented with 10% FBS and 1% penicillin–streptomycin. MCF10A cells were grown in DMEM/F-12 supplemented with 5% horse serum, 10 µg/mL insulin, 20 ng/mL epidermal growth factor (EGF), 100 ng/mL hydrocortisone, and 100 ng/mL cholera toxin. All cell lines were authenticated by short tandem repeat (STR) profiling and routinely tested to confirm absence of mycoplasma contamination.

### Plasmid and transfections

GFP-mROCK1 (Addgene plasmid # 187296; http://n2t.net/addgene:187296; RRID:Addgene_187296) and GFP-mROCK1-delta3 (Addgene plasmid # 101291; http://n2t.net/addgene:101291; RRID:Addgene_101291) were a gift from Alpha Yap^20^. The pCMV3-PLS3-GFPSpark plasmid (Cat. HG15042-ACG) was purchased from Sino Biological (Beijing, China). The pC3-EGFP plasmid was kindly provided by the laboratory of Thomas Kaufmann (University of Bern).

Transient transfection was performed using Lipofectamine™ 2000 according to the manufacturer’s instructions. Cells were incubated for 4 h in serum-free Opti-MEM containing 1 µg of plasmid DNA per well and Lipofectamine™ 2000 at the following volumes: 5 µL per well (6-well plates), 2 µL per well (24-well plates), and 3 µL per well (WillCo dishes). After transfection, the medium was replaced with complete growth medium, and cells were used for experiments 24 h later.

### Immunofluorescence

MCF7 cells were seeded at 60,000 cells/well on 13-mm glass coverslips and cultured overnight. The next day, cells were treated with 1 µM narciclasine for 24 h for E-cadherin and PLS3 immunofluorescence, or for 6 h for pMLC2 staining. After treatment, cells were fixed with 4% paraformaldehyde for 10 min at room temperature and permeabilized with 95% ethanol containing 5% glacial acetic acid. For E-cadherin/LAMP1 staining, cells were incubated overnight at 4 °C with rabbit anti-E-cadherin (24E10, Cell Signaling Technology, #3195S) and mouse anti-LAMP1 (BD Pharmingen, #555798), followed by Alexa Fluor® 488–conjugated anti-rabbit and Alexa Fluor® 555–conjugated anti-mouse secondary antibodies (Thermo Fisher Scientific) for 2 h at room temperature. For PLS3 staining, cells were incubated with mouse anti-PLS3 (Invitrogen, MA5-27772) followed by Alexa Fluor® 555–conjugated anti-mouse secondary antibody (Thermo Fisher Scientific). For pMLC2/actin staining, cells were incubated with rabbit anti-phospho-MLC2 (Ser19, Cell Signaling Technology, #3671S) and mouse anti-actin (Merck, A5441), followed by Alexa Fluor® 488–conjugated anti-rabbit and Alexa Fluor® 555–conjugated anti-mouse secondary antibodies (Thermo Fisher Scientific). Coverslips were mounted on microscope slides using ProLong® Diamond Antifade Mountant containing DAPI (Thermo Fisher Scientific). Images were acquired on a Leica SP8 confocal laser-scanning microscope using an HC PL APO CS2 40×/1.30 oil immersion objective for PLS3 imaging and an HC PL APO CS2 63×/1.40 oil immersion objective for pMLC2 imaging. Excitation wavelengths were 405 nm (DAPI), 488 nm (Alexa Fluor 488), and 552 nm (Alexa Fluor 555), and emission was collected using appropriate spectral detection windows. For each field of view, z-stacks spanning the entire cell volume were acquired with 0.6 µm step size, a resolution of 1024 × 1024 pixels, and a pinhole set to 1 Airy unit. Image stacks were processed in ImageJ (Fiji). To quantify the PLS3 membrane-to-cytosol ratio, a custom CellProfiler pipeline was used in which nuclei were segmented with the IdentifyPrimaryObjects module using adaptive thresholding (Otsu), cell boundaries were defined with IdentifySecondaryObjects on the PLS3 channel using a propagation method with a global threshold based on minimum cross-entropy. The cytoplasmic compartment was defined by subtracting the nuclear objects from the total PLS3-positive cell area until the boundary of the nuclei was reached. The membrane object was then generated using the cytoplasm as input and applying the operation “shrink object to a point” to define the peripheral compartment. The *RelateObjects* module was used to assign parent–child relationships between membrane objects (parent) and nuclei (child). Mean fluorescence intensities in the cytoplasmic and membrane compartments were quantified from the PLS3 channel using the *MeasureObjectIntensity* module. Finally, the *CalculateMath* module was used to compute the membrane-to-cytosol intensity ratio for each cell.

### Entosis detection and quantification

Entosis was detected by immunofluorescence staining for β-catenin and LAMP1. For experiments shown in Fig. 2A, 60,000 MCF7 cells/well were seeded onto 13-mm round glass coverslips placed in 24-well plates and allowed to adhere overnight. Cells were then treated with vehicle (DMSO) or narciclasine (0.1, 0.5, or 1 µM) for 24 h. For experiments shown in Fig. 4, MCF7 cells were pre-treated with 20 µM Y27632 for 1 h prior to the addition of 1 µM narciclasine (for 24 h). Control conditions included vehicle alone, narciclasine alone, and Y27632 alone. The same treatment conditions were applied to EFM19 and ZR75-1 cells. For experiments shown in Fig. 5E, MCF7 cells were treated with vehicle (DMSO) or 1 µM narciclasine for 6 h. For experiments shown in Fig. 6, MCF7 cells were pretreated with 50 µM zVAD-fmk for one hour followed by treatment with 1 µM narciclasine (24 h). Vehicle-treated cells were used as controls. Following drug incubation, cells were washed with PBS and fixed in 4% paraformaldehyde (PFA) for 10 min at room temperature. Cells were then permeabilized with 95% ethanol/5% glacial acetic acid for 10 min and blocked in 5% BSA in PBS. Primary antibodies against rabbit β-catenin (Epredia, RB-9035-P1, 1:200) and mouse LAMP1 (BD Pharmingen, 555798, 1:200) were applied overnight at 4 °C, followed by Alexa Fluor® 488-conjugated anti-rabbit (A11008, 1:500; Thermo Fisher Scientific) and Alexa Fluor® 555-conjugated anti-mouse secondary antibodies (A21424, 1:500; Thermo Fisher Scientific). Coverslips were mounted using ProLong® Diamond Antifade Mountant with DAPI (Thermo Fisher Scientific) to visualize nuclei. Images were acquired using a Leica SP8 confocal laser-scanning microscope equipped with an HC PL APO CS2 40×/1.30 oil immersion objective. Excitation was performed using 405 nm, 488 nm, and 552 nm lasers for DAPI, Alexa Fluor 488, and Alexa Fluor 555, respectively. Emission detection ranges were set to 410–483 nm (DAPI), 493–556 nm (Alexa Fluor 488), and 557–729 nm (Alexa Fluor 555). Z-stacks were acquired across the entire cell volume with an optical section thickness of 0.6 µm at a resolution of 1024 × 1024 pixels and a scan speed of 600 Hz. Entotic events were quantified using 3D confocal z-stacks. For each condition, at least 500–1,000 cells per sample were analyzed across 5–7 randomly selected fields of view (FOV). Entotic events were quantified based on the identification of round-shaped internal cells (Hoechst-, LAMP1- and β-catenin–positive) enclosed within a large vacuole inside a host cell. The host cell was defined as Hoechst-, β-catenin- and LAMP1-positive, displaying punctate or ring-like LAMP1 structures and a characteristic crescent-shaped nuclear morphology. Both live and dead inner cells, as well as structures containing multiple internalized cells, were counted as a single entotic event. The number of entotic structures was normalized to the total number of nuclei per field of view and expressed as a percentage.

### Volumentric Electron Microscopy (SEM Array Tomography)

For electron microscopy studies, 100,000 MCF7 cells per well were seeded in a Lab-Tek chamber slides of 4 wells (Nalge Nunc International, Naperville, IL) and treated with 1uM narciclasine for 24 hours. After incubation, cells were fixed in 3 % glutaraldehyde in 0.1 M phosphate buffer (PB) for 1 hour at 37°C. Last, were washed in 0.1 M phosphate buffer (PB) for 5 times and stored at 4 °C. The samples were postfixed in 2% OsO4 for 1 hour at room temperature and stained in 2% uranyl acetate in the dark for 2 h at 4 °C. Then, were rinsed in destilled water, dehydrated in ethanol and infiltrated overnight in Durcupan resin (Sigma-Aldrich, St. Louis, USA). Following polymerization, embedded cultures were detached from the wells and glued to Durcupan blocks. An asymmetric blockface was created with parallel leading and trailing edges to facilitate the formation of straight ribbons during sectioning. Serial ultrathin sections (approx. 70–90 nm thick) were cut using an ultramicrotome Leica UC-6 (Leica microsystems, Wetzlar, Germany), equipped with an Ultra Jumbo diamond knife (Diatome Ltd, Nidau, Switzerland) and collected with a custom water withdrawal device. Sequential ribbons were then treated with xylene vapor to minimize compression and stretch the sections. The entire array was transferred onto an Indium Tin Oxide (ITO)-coated glass coverslip and to enhance ultrastructural contrast for electron microscopy, the sections were post-stained with lead citrate (Reynold’s solution). The dry coverslip was mounted onto an SEM stub using conductive copper tape to ensure electrical grounding. Images were acquired using a FESEM (Field Emission Scanning Electron Microscope) equipped with a sensitive backscatter electron (BSE) detector (GeminiSEM 460 with BSD). The entire section array was mapped at low resolution to locate all sections. ROIs were then defined on a single section and propagated across the series using automated array tomography software (ATLAS 5). Automated functions for focus, stigmation, and stage movement were employed to capture the 3D dataset at the required resolution, with a final acquisition pixel size of 5 nm per pixel. The resulting 2D image series was aligned into a 3D volume stack. Initial alignment utilized rigid transformations (rotation and XY translation) to correct for physical distortions introduced during sectioning and collection. Final registration was performed using cross-correlation or Scale Invariant Feature Transform (SIFT) algorithms within software packages such as Fiji (TrakEM2), IMOD, or Amira. The aligned volumes were validated by inspecting XZ and YZ projections to ensure ultrastructural continuity. Following alignment and initial segmentation, the processed 3D volumes were imported into Webknossos (high-throughput 3D data analysis platform). This software was utilized for the final visualization, manual proofreading, and analysis of the ultrastructural features.

### Proteomics and data analysis

Cells were lysed in RIPA buffer (50 mM Tris-HCl, pH 8.0, 150 mM NaCl, 1% NP-40 substitute, 0.5% sodium deoxycholate and 0.1% SDS). A total of 10 µg of protein in 20 µL was prepared for further processing. Samples were adjusted to 30 µL with 50 mM ammonium bicarbonate (ABC). Disulfide bonds were reduced by incubating samples for 20 min at 60°C in 35 µL of 2 mM DTT prepared in 50 mM ABC. Samples were allowed to cool to room temperature, and free sulfhydryl groups were alkylated in 40 µL of 5.5 mM iodoacetamide (IAM) for 30 min at room temperature in the dark. Alkylated samples were processed using the SP3 protocol with minor modifications^21,22^. Proteins were incubated with SP3 resin. Then, 20 µL of beads were used to achieve a 1:20 protein-to-beads ratio, and acetonitrile (ACN) was added to a final concentration of 70%. After bead-based cleanup, proteins were digested with trypsin (500 ng in 100 µL of 50 mM ABC) at 37°C overnight. Digested peptides were acidified with 10% TFA to a final concentration of 1%. The final sample volume was 110 µL, corresponding to a theoretical peptide concentration of 181 ng/µL. Approximately 200 ng of each digest were diluted to 20 µL with 0.1% formic acid (FA) and loaded onto Evotip Pure tips (EvoSep) according to the manufacturer’s instructions. Sample injection order was fully randomized using HyStar (Bruker). LC–MS/MS analysis was performed on a timsTOF fleX mass spectrometer (Bruker). Peptides loaded on Evotip Pure tips were eluted onto an analytical column (Endurance 15 cm × 150 µm, 1.5 µm; Evosep) using the Evosep One system and separated using the manufacturer-defined 30 SPD chromatographic method. Eluted peptides were ionized using a CaptiveSpray source at 1600 V and 180°C and analyzed in diaPASEF mode (method: diaPASEF longGradient) with the following settings: TIMS settings: custom mode; 1/K0 range 0.6–1.6 V·s/cm²; ramp time 100 ms; duty cycle 100%; ramp rate 9.42 Hz; MS averaging 1; auto calibration off. MS settings: scan range 100–1700 m/z; ion polarity positive; scan mode diaPASEF.

System performance was monitored using 50 ng of HeLa digest, resulting in identification of 7,417 proteins using diaPASEF and the 30 SPD gradient. Raw diaPASEF data were processed using DIA-NN v1.8 via FragPipe 21.1. An in silico–predicted spectral library was generated from the SwissProt Human database (2024-07-23 release). Quantification results were exported as Excel files containing unique genes and protein groups filtered at FDR ≤ 1%.

Differential expression analysis was performed using FragPipe-Analyst (http://fragpipe-analyst.nesvilab.org/)^23^. Raw DIA-NN output (diann-output.pg_matrix.tsv) was filtered to remove contaminants, proteins identified by a single peptide, and proteins not consistently quantified within the same condition. Data were normalized under the assumption that the majority of proteins do not change between conditions. Intensities were log2-transformed, samples grouped by condition, and missing values imputed using a left-shifted Gaussian distribution (1.8 s.d. downshift, width 0.3). Differential expression was performed using protein-wise linear models with empirical Bayes moderation implemented in limma (R/Bioconductor)^24^. Proteins with FDR < 0.05 and |log2 fold change| > 1 were considered significantly regulated. Differential expression results for narciclasine- versus vehicle-treated cells were analyzed from an exported table containing gene symbols, log2 fold changes and adjusted p-values. Gene identifiers were cleaned by splitting multi-symbol entries and retaining unique symbols. Proteins were considered significantly regulated using thresholds of |log2 fold change| > 1 and FDR < 0.05. Volcano plots were generated in R using ggplot2, displaying log2 fold change versus −log10(FDR). The most significant upregulated and downregulated proteins (top 10 per group by FDR) were annotated. Over-representation analysis of Gene Ontology (GO) Biological Process terms was performed using clusterProfiler^25^ with human annotations from org.Hs.eg.db. Significantly regulated proteins (FDR < 0.05, |log2FC| > 1) were mapped to Entrez identifiers and tested for enrichment using Benjamini–Hochberg correction. The top enriched biological processes were ranked by adjusted p-value and visualized as bar plots. To assess pathway-level shifts independently of arbitrary significance cutoffs, preranked gene set enrichment analysis was performed using fgsea. All quantified proteins were ranked by log2 fold change. Reactome pathway gene sets (MSigDB C2:CP:REACTOME collection) were retrieved using msigdbr and tested with 10,000 permutations. Pathways with FDR < 0.05 were considered significantly enriched. Cytoskeleton- and mechanotransduction-related pathways were highlighted based on curated keyword filtering.

### Time-lapse imaging

For experiments shown in Fig. 6C, MCF7 cells were seeded at a density of 30,000 cells per well in WillCo-dish® glass-bottom dishes (22-mm aperture) fitted with 4-well Culture-Inserts (Ibidi). After overnight attachment, cells were incubated with 0.2 µM LysoTracker™ Deep Red and 0.5 µg/mL Hoechst 33342 for 30 min at 37°C. Cells were then treated with vehicle (DMSO), 1 µM narciclasine, or 1 µM narciclasine plus 50 µM zVAD-fmk. Time-lapse imaging was performed using a Leica DMI6000B inverted microscope equipped with an HC PL FLUOTAR 20×/0.50 NA dry objective and an HBO 100 mercury short-arc lamp. Cells were maintained throughout imaging at 37°C in a humidified environmental chamber with 5% CO₂.

Images were acquired at 1392 × 1040 pixel resolution with 1 × 1 binning every 10 min for 24 h. Hoechst was imaged using excitation filter BP 340–380 nm and emission filter LP 425 nm (exposure time: 262 ms). LysoTracker™ Deep Red was detected using excitation filter BP 520/60 nm and emission filter BP 700/75 nm (exposure time: 300 ms). Time-lapse recordings were analyzed using ImageJ. Internalized cells were manually classified into the following categories: no change (internal cell stably retained within the engulfing cell at 24 h), death (defined by accumulation of LysoTracker signal within the internal cell), division, or escape. Event distribution was calculated as the percentage of each category normalized to the total number of entotic events. For experiments shown in Fig. 2C, MCF7 cells were seeded at 100,000 cells per WillCo-dish®. The following day, cells were incubated with 0.5 µg/mL Hoechst 33342 for 30 min in FluoroBrite™ DMEM, treated with 1 µM narciclasine, and imaged using an HCX PL FLUOTAR L 40×/0.60 NA dry objective on the same Leica AF6000 LX system under identical environmental conditions (37°C, 5% CO₂). Hoechst acquisition settings were identical to those described above.

For PLS3–GFP time-lapse experiments, images were acquired every 5 min. GFP was detected using excitation filter BP 470/40 nm, whereas Hoechst was imaged using the same acquisition settings described above. Imaging was performed under identical environmental conditions (37°C, 5% CO₂).

### Live-cell imaging

MCF7 cells were seeded on WillCo-dish® glass-bottom dishes (22-mm aperture) at a density of 100,000 cells per dish and allowed to attach overnight. Cells were treated with either vehicle (DMSO) or 1 µM narciclasine for 24 h. Following treatment, cells were incubated for 30 min at 37 °C in FluoroBrite™ DMEM containing WGA Deep Red, 0.2 µM LysoTracker Red DND-99, and 0.5 µg/mL Hoechst. Dishes were then transferred to a Leica SP8 confocal laser-scanning microscope equipped with an HC PL APO CS2 63×/1.40 oil immersion objective, maintaining environmental conditions at 37 °C and 5% CO₂ during imaging. Excitation and emission settings were as follows: 405 nm for Hoechst (410–547 nm emission), 552 nm for LysoTracker (557–644 nm emission), and 638 nm for WGA Deep Red (644–784 nm emission). Z-stacks spanning the full cellular volume were acquired at 0.6 µm intervals with a resolution of 1024 × 1024 pixels and a pinhole set to 1 Airy unit. Image stacks were processed using ImageJ (Fiji). Early-stage entotic cells were defined as Hoechst-positive internalized cells displaying punctate LysoTracker signal, whereas late-stage entotic cells were defined as Hoechst-positive internalized cells exhibiting a rounded accumulation of LysoTracker within the entotic vacuole. The percentage distribution of early and late stages was calculated by dividing the number of early- or late-stage entotic cells by the total number of entotic cells.

### Correlative Light and Electron Microscopy (CLEM)

For correlative light and electron microscopy (CLEM) analysis, MCF7 cells were seeded at a density of 100,000 cells per well in 4-well Permanox Lab-Tek chamber slides (Nalge Nunc International) and allowed to attach overnight. The following day, cells were treated with 1 µM narciclasine for 24 hours. Cells were then stained with 0.5 µg/mL Hoechst and 0.2 µM LysoTracker for 30 minutes at 37°C. Cells were fixed in 3% glutaraldehyde in 0.1 M phosphate buffer (PB) for 20 minutes at 37°C and subsequently for 2 hours at room temperature. Slides were then washed five times in 0.1 M PB. Samples were mounted using Fluoromount aqueous mounting medium and covered with a larger coverslip. Events were first identified using a Leica DM6000 fluorescence microscope, and their positions were marked using a Nikon microscope object marker. These positions were subsequently retrieved, and confocal images were acquired using a Leica TCS SP8 inverted laser scanning confocal microscope equipped with an HC PL APO CS2 63×/1.40 oil immersion objective, as described above. Excitation wavelengths were 405 nm for Hoechst and 552 nm for LysoTracker Red. Optical sections were acquired every 0.6 µm at a resolution of 1024 × 1024 pixels using an Airy 1 pinhole diameter.

After fluorescence imaging, the positions were further marked using an Olympus IX71 laser microdissection microscope equipped with P.A.L.M. technology, in cut mode. Slides containing the selected events were then processed for transmission electron microscopy (TEM). Samples were post-fixed in 2% OsO₄ for 1 hour at room temperature and stained with 2% uranyl acetate in the dark for 2 hours at 4°C. Samples were rinsed in distilled water, dehydrated in ethanol, and infiltrated overnight with Durcupan resin (Sigma-Aldrich). Following polymerization, embedded cultures were detached from the wells and glued to Durcupan blocks. Ultrathin sections (0.08 µm) were cut using an Ultracut UC-6 (Leica Microsystems), stained with lead citrate (Reynolds solution), and examined using a FEI Tecnai Spirit BioTwin transmission electron microscope (Thermo Fisher Scientific™). Images were acquired using Radius software (version 2.1) with a Xarosa digital camera (EMSIS GmbH). Finally, confocal images were rotated to match the corresponding TEM images, and correlation between fluorescence and electron microscopy images was performed using ICY software with the Ec-CLEM plugin employing non-rigid deformation^26^.

### Immunoblotting

Whole-cell lysates were prepared in RIPA buffer (50 mM Tris-HCl, pH 8.0, 150 mM NaCl, 1% NP-40 substitute, 0.5% sodium deoxycholate and 0.1% SDS) supplemented with PMSF, pepstatin A and leupeptin for standard immunoblotting. For detection of phosphorylated proteins, lysates were prepared using PhosphoSafe™ Extraction Reagent according to the manufacturer’s instructions. Protein samples were resolved by SDS–PAGE and transferred onto nitrocellulose membranes using standard procedures. Membranes were blocked in 5% non-fat milk in TBS-T or in 5% BSA in TBS-T when detecting phospho-specific antibodies and incubated with primary antibodies against ROCK1 (rabbit monoclonal, 1:1000; Cell Signaling Technology, #28999S), RhoA (rabbit monoclonal, 1:1000; Cell Signaling Technology, #2117S), PLS2 (rabbit monoclonal, 1:1000; Atlas Antibodies, #HPA019493), PLS3 (mouse monoclonal, 1:1000; Invitrogen, #MA5-27772), α-tubulin (rabbit polyclonal, 1:3000; Cell Signaling Technology, #2144S), phospho-MLC2 (Ser19) (rabbit, 1:1000; Cell Signaling Technology, #3671S), and GFP (chicken polyclonal, 1:1000; Rockland Immunochemicals, #600-901-215). After washing, membranes were incubated with horseradish peroxidase (HRP)-conjugated secondary antibodies, including goat anti-mouse IgG-HRP (Merck, #A4416), goat anti-rabbit IgG-HRP (Merck, #A6154), and goat anti-chicken H&L-HRP (Rockland Immunochemicals, #603-103-002). Immunoreactive bands were detected using enhanced chemiluminescence (ECL) reagents (Cytiva, #RPN2209, or NZYtech, #MB40201) and imaged using an Amersham ImageQuant™ 800 system (Cytiva).

### Gene silencing

Transient silencing of ROCK1 and PLS3 was performed by RNA interference. A non-targeting control siRNA (Santa Cruz Biotechnology, #sc-44237) was used as a negative control. Cells were transfected using Lipofectamine™ 2000 (Thermo Fisher Scientific) according to the manufacturer’s instructions. Briefly, cells were incubated for 4 h in serum-free Opti-MEM™ containing siRNA and Lipofectamine™ 2000. ROCK1 was silenced using ON-TARGETplus SMARTpool siRNA (Dharmacon) at a final concentration of 500 nM, and PLS3 was silenced using siRNA (Santa Cruz Biotechnology, #sc-43215) at a final concentration of 200 nM. Lipofectamine™ 2000 was used at 5 µL per well in 6-well plates and 2 µL per well in 24-well plates. After transfection, medium was replaced with complete growth medium.

For ROCK1 silencing, protein lysates were collected 24 h post-transfection from 6-well plates for immunoblot analysis. In parallel, cells seeded in 24-well plates were treated with vehicle or 1 µM narciclasine and subjected to entosis quantification as described above. For PLS3 silencing, protein lysates were collected 48 h post-transfection, and entosis assays were performed under the same treatment conditions.

### Survival analysis

Survival analyses were performed using publicly available breast cancer transcriptomic datasets interrogated through an online Kaplan–Meier survival analysis platform. Gene expression of PLS3 (T-plastin) was assessed using the Affymetrix probe set 201215_at. Relapse-free survival (RFS) was selected as the clinical endpoint. Patients were stratified into high and low expression groups using the median expression value within the analyzed cohort (cutoff value: 2852; expression range: 17–17,790). Median-based dichotomization was applied within the selected dataset, and no optimized cutpoint selection was used. Follow-up time was not truncated, and patients were censored at the follow-up threshold when applicable. All available breast cancer cohorts were included without restriction by ER status (IHC or array), PR status, HER2 status, molecular subtype (St Gallen or PAM50), lymph node status, tumor grade, TP53 status, or Pietenpol subtype. No cohort-specific filtering was applied. Quality control measures included removal of redundant samples and exclusion of biased arrays according to platform-defined quality control criteria. The proportional hazards assumption was tested and confirmed.

### Presentation of data and statistical analysis

All experiments were performed in at least three independent biological replicates. Representative images are shown where applicable. In most cases, data are presented as percentages of absolute values. For scatter plots with bar overlays, data are shown as mean ± SEM; for violin plots, data are presented as median ± interquartile range (IQR). Statistical analyses were performed using GraphPad Prism v10 (GraphPad Software, San Diego, CA, USA). Comparisons between two groups were conducted using a paired or unpaired two-tailed Student’s t-test. Multiple-group comparisons were analysed using one-way or two-way analysis of variance (ANOVA), followed by Bonferroni’s post hoc test where appropriate. Proteomics data were analysed using RStudio. A p value < 0.05 was considered statistically significant.

## Data availability

The mass spectrometry proteomics data have been deposited to the ProteomeXchange Consortium via the PRIDE^27^ partner repository with the dataset identifier PXD075153. Data supporting this work are available in the Article. All other data supporting the findings of this study, including imaging files, are available from the corresponding author upon reasonable request.

## Code availability

The proteomics analysis code and fully reproducible computational environment are publicly available in Zenodo at DOI: 10.5281/zenodo.18847057. The repository includes the differential expression results table (DE_results.csv), analysis scripts, generated figures and tables, and a renv lockfile enabling exact restoration of the R package environment. All results can be reproduced by restoring the environment with renv and executing the main analysis script provided in the repository.

## RESULTS

### Constitutive ROCK1 activation triggers entotic invasion

First, we sought to determine whether active ROCK1 is sufficient to induce entosis. To this end, MCF7 cells were transfected with either a mouse GFP-ROCK1 construct or GFP-mROCK1-Δ3, a constitutively active ROCK1 mutant lacking the C-terminal autoinhibitory region (Figure 1A and Supplementary Figure 1A). Western blot analysis revealed ROCK1 bands at distinct molecular weights corresponding to the GFP tag and the truncated size of the Δ3 construct, which migrates faster due to the deletion of regulatory domains (Figure 1B and Supplementary Figure 1B). We also evaluated the phosphorylation of MLC2 (pMLC2), a downstream readout of ROCK1 activity. Consistent with constitutive activation, cells expressing ROCK1 Δ3 displayed increased pMLC2 levels compared to controls (Figure 1B), after 6 hours transfection. We next performed live-cell imaging of MCF7 cells transfected with either GFP–ROCK1 or GFP–ROCK1 Δ3 to determine whether enforced ROCK1 activation is sufficient to induce entotic structures formation. Analyses were restricted to GFP-positive cells, and both the overall induction and quantification of CIC structures were assessed (Figure 1C and D). In both conditions, we detected CIC structures displaying canonical morphological hallmarks of entosis, including the characteristic crescent-shaped nucleus of the external cell (EC) and complete internalization of a rounded inner cell (IC) within an actin-positive EC (Supplementary Figure 1C and D; Figure 1C). In addition, the presence of an entotic vacuole was evident as a pericellular shadow surrounding the internalized cell in fluorescence images and as an optically empty space in phase-contrast images (Supplementary Figure 1C and D). Quantification of entotic events among GFP-positive cells revealed a significant increase in entosis frequency in cells expressing GFP-ROCK1 Δ3 compared with GFP-ROCK1 (Figure 1D). We further classified entotic events based on the GFP status of the internal and external cells, identifying three populations: GFP-positive cells internalized into GFP-negative cells (Internal), GFP-negative cells internalized into GFP-positive cells (External), and CIC structures in which both cells were GFP-positive (Both). Analysis of the distribution of these populations revealed that GFP-ROCK1 Δ3 expression was associated with a higher proportion of GFP-positive cells acting as the internal cell, whereas GFP-ROCK1 expression preferentially resulted in CIC structures in which both cells were GFP-positive (Figure 1E).

**Figure 1.**
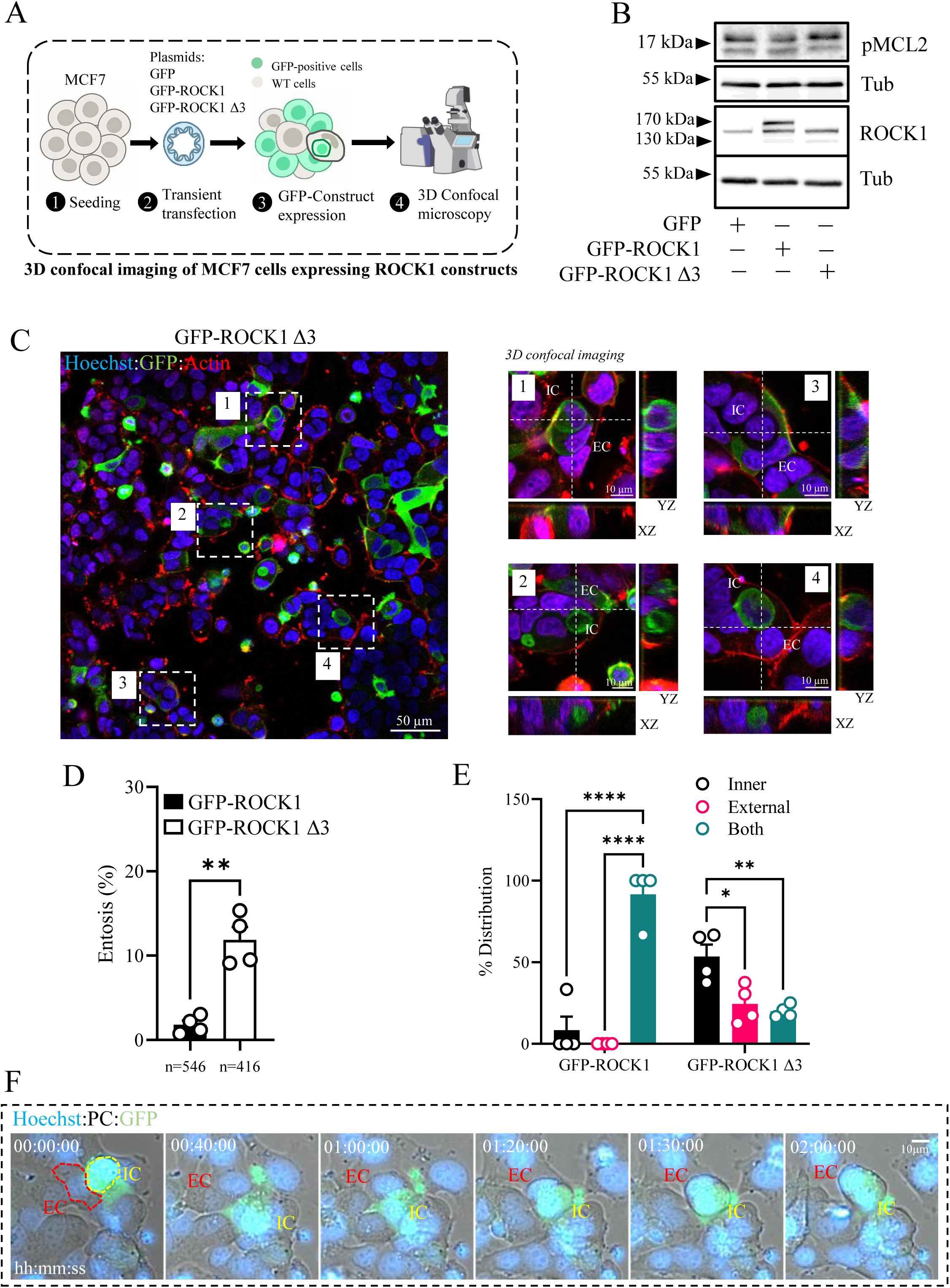
Constitutively active ROCK1 but not its abundance induces CIC formation. **(A)** Schematic overview of ROCK1 construct transfection used to assess entosis induction. **(B)** Representative immunoblots showing ROCK1, phosphorylated MLC2 (pMLC2), and tubulin (loading control) in MCF7 cells transfected with GFP, GFP–ROCK1, or constitutively active GFP–ROCK1 Δ3. Immunoblot analysis of pMLC2 was performed 6 h post-transfection, whereas ROCK1 expression was assessed at 24 h. **(C)** Representative 3D confocal images of live MCF7 cells expressing GFP–ROCK1 Δ3 and stained with SiR-Actin (red) and Hoechst (blue). White arrows indicate entotic structures. Boxed regions showing representative entotic structures are magnified, with corresponding orthogonal z-stack views shown alongside each image. **(D)** Quantification of entotic events in MCF7 cells expressing GFP–ROCK1 or GFP–ROCK1 Δ3. **(E)** Distribution percentages of GFP-positive cells participating in CIC structures as outer cells, inner cells, or both. **(F)** Representative time-lapse imaging (phase contrast, Hoechst, and GFP) capturing an entotic event in MCF7 cells expressing GFP–ROCK1 Δ3. Yellow dotted lines outline inner cells, and red dotted lines outline host cells. Experiments were performed in biological triplicate (n = 3). Data are presented as mean ± SEM. Statistical analysis in panel D was performed using a paired two-tailed Student’s t-test (**p < 0.01), whereas panel E was analyzed using two-way ANOVA followed by Bonferroni post hoc test (*p < 0.05, **p < 0.01, ****p < 0.0001).

Finally, induction of entosis was directly confirmed by time-lapse imaging. As shown in Figure 1F, MCF7 cells expressing GFP-ROCK1-Δ3 displayed the typical morphological sequence of entosis, including cell rounding, polarized invasion, and progressive internalization.

### Narciclasine induces entosis in MCF7 cells

Next, we searched the literature for selective ROCK1 activators and identified narciclasine, which has been reported to activate RhoA/ROCK1 signaling^28^. MCF7 cells were treated with increasing concentrations of narciclasine (0.1 µM, 0.5 µM, and 1 µM) for 24 hours, followed by immunofluorescence staining for LAMP1 and β-catenin and acquisition of Z-stack images by confocal microscopy. Strikingly, narciclasine treatment induced multiple types of CIC structures at all tested concentrations, consistent with entotic events (Figure 2A). First, we observed structures characterized by accumulation of LAMP1 around the IC, forming a ring-like pattern indicative of an entotic vacuole. Second, in agreement with previous reports^16,29^, early-stage entotic structures exhibited β-catenin localization at the inner cell interface, while in late stage entotic structures β-catenin was not detected due to IC degradation. Finally, we detected entotic structures displaying a characteristic crescent-shaped (“half-moon”) nucleus in the EC, with the IC enclosed within a large LAMP1-positive entotic vacuole, consistent with more advanced entotic stages (Fig. 2A). Based on the characteristic features described above, we next quantified entotic events and observed a dose-dependent increase in their frequency across narciclasine concentrations, ranging from approximately 10% at 0.1 µM to ∼15% at 1 µM (Figure 2B). To further validate that these structures reflected bona fide entotic invasion, we performed time-lapse imaging. As shown in Figure 2C and Video 1, MCF7 cells treated with 1 µM narciclasine exhibited dynamic cellular behaviors and morphological changes consistent with active entotic internalization of one cell into another. Additional imaging using Hoechst nuclear staining, LysoTracker, and phase-contrast microscopy revealed distinct fates of entotic structures following narciclasine treatment. We observed cellular behaviors consistent with active invasion (Supplementary Figure 2A and Video 2), invasion followed by subsequent escape of the internalized cell (Supplementary Figure 2B and Video 3), and internalized cells undergoing entotic cell death, as indicated by LysoTracker accumulation (Supplementary Figure 2C and Video 4).

**Figure 2.**
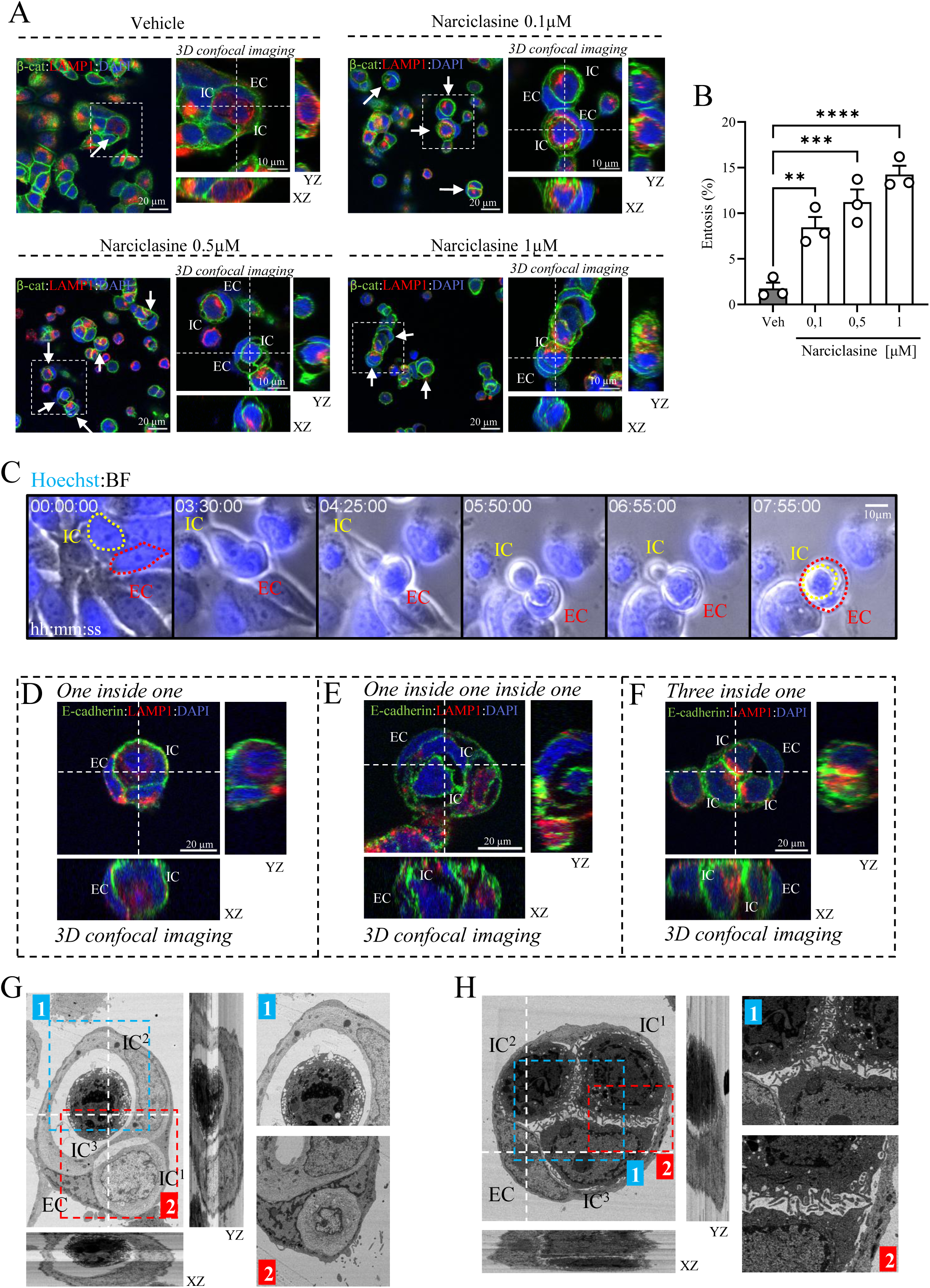
Narciclasine induces entosis in MCF7 cells. **(A)** Representative 3D confocal images of vehicle- and narciclasine-treated cells. MCF7 cells were treated with increasing concentrations of narciclasine (0.1, 0.5, and 1 µM) and vehicle (DMSO) for 24 h. Entotic structures were visualized by immunofluorescence staining for β-catenin (membrane, green), LAMP1 (lysosomal structures, red) and DAPI (nuclei, blue) followed by z-stack confocal microscopy. White arrows indicate entotic structures. Selected regions (boxed) showing representative entotic structures are magnified, with corresponding orthogonal z-stack views shown adjacent to each image. **(B)** Summary of quantification of entotic events in MCF7 cells treated with vehicle or increasing concentrations of narciclasine. **(C)** Representative time-lapse images (phase contrast and Hoechst) showing induction of entosis in MCF7 cells treated with 1 µM narciclasine. Yellow dotted lines outline inner cells, and red dotted lines outline host cells. **(D)** Representative 3D confocal image of a one-cell-inside-another entotic structure in MCF7 cells treated with 1 µM narciclasine, visualized by immunofluorescence staining for E-cadherin (green), LAMP1 (red), and DAPI (blue). **(E)** Representative confocal image of a nested entotic configuration (cell-in-cell-in-cell) in MCF7 cells treated with 1 µM narciclasine, visualized by E-cadherin, LAMP1, and DAPI staining. **(F)** Representative confocal image of three cells internalized within a single host cell following treatment with 1 µM narciclasine, visualized by E-cadherin, LAMP1, and DAPI staining. **(G)** Volumetric electron microscopy image of a nested entotic structure in MCF7 cells treated with 1 µM narciclasine. **(H)** Volumetric electron microscopy image of three cells inside one in MCF7 cells treated with 1 µM narciclasine. For all confocal and electron microscopy panels, orthogonal z-stack views and magnified insets are provided. Experiments were performed in biological triplicate (n = 3). Data are presented as mean ± SEM and were analyzed by one-way ANOVA followed by Bonferroni post-hoc test (**p < 0.01, ***p < 0.001, ****p < 0.0001).

We also performed immunofluorescence staining to detect LAMP1 and E-cadherin expression in entotic structures. Again, we visualized several types of entotic configurations: a classical one-inside-one structure at a late stage, in which E-cadherin expression is abolished (Fig. 2D and Supplementary Fig. 2D); a one-inside-one-inside-one structure at an early stage, where E-cadherin is present at the inner cell forming a ring-like pattern (Fig. 2E and Supplementary Fig. 2E); and a three-cells-inside-one configuration, again showing E-cadherin enrichment at cell–cell contact sites (Fig. 2F and Supplementary Fig. 2F).

To further confirm the presence of entotic structures and to visualize their ultrastructural organization, we employed volumetric electron microscopy (vEM). Consistent with our light microscopy observations, treatment with 1 µM narciclasine robustly induced CIC (Supplementary Figure 3A). Notably, vEM revealed complex entotic architectures, including nested CIC configurations in which two cells were internalized within a single EC, with one internalized cell itself containing another cell (Figure 2G, Video 5). In addition, we observed more complex CIC configurations involving multiple invasion events, resulting in structures containing three cells within a single EC (Figure 2H, Video 6). We also observed large entotic vacuoles containing a single dead internalized cell, consistent with late-stage entotic degradation (Supplementary Figure 3B), together with simpler, canonical one-cell-inside-one-cell structures (Supplementary Figure 3C).

Next, we performed a series of experiments to further characterize and confirm entotic events in control and narciclasine-treated cells. We first conducted live-cell imaging to examine morphological features of early- and late-stage entotic structures using plasma membrane staining (WGA), nuclear labeling (Hoechst), and lysosomal accumulation (LysoTracker). Confocal Z-stack imaging confirmed complete internalization of IC within EC in both conditions (Fig. 3A and B). Consistent with previous reports^19^, these markers enabled discrimination of early and late entotic stages, with punctate LysoTracker signal in early IC and EC and strong lysosomal accumulation in late-stage structures inside the entotic vacuole (Fig. 3A and B). Quantification of population distribution revealed a predominance of LysoTracker-negative IC in vehicle-treated cultures, whereas narciclasine treatment resulted in comparable proportions of LysoTracker-negative and LysoTracker-positive IC (Fig. 3C).

**Figure 3.**
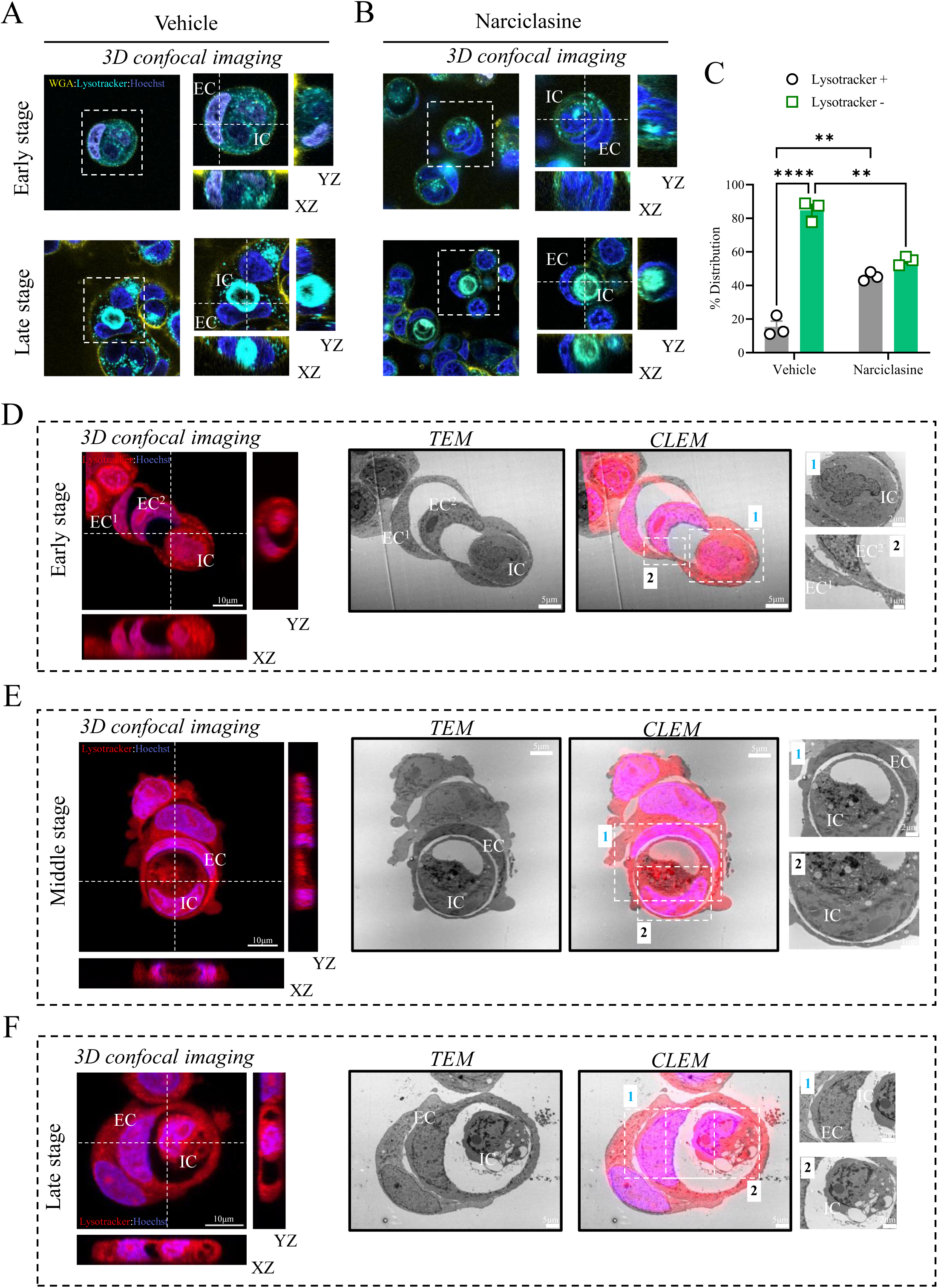
Characterization of entotic structures induced by narciclasine. **(A–B)** MCF7 cells were treated with vehicle (DMSO) or 1 µM narciclasine for 24 h. Entotic structures were visualized by 3D live-cell confocal imaging following staining with WGA (plasma membrane, yellow), LysoTracker Red (lysosomes, cyan), and Hoechst (nuclei, blue). Representative early- and late-stage entotic structures are shown. **(C)** Summary of quantification of inner-cell stage distribution based on lysosomal accumulation. Late-stage inner cells were defined by round LysoTracker-positive accumulation, whereas early-stage inner cells displayed punctate or absent LysoTracker signal. **(D–F)** Correlative light–electron microscopy (CLEM) analysis of entotic ultrastructure in MCF7 cells treated with 1 µM narciclasine. Representative confocal images of Hoechst (blue) and LysoTracker (red) identify early (D), middle (E), and late (F) entotic stages (right panels). Corresponding transmission electron microscopy (TEM) images are shown in the center, and merged correlative overlays are shown on the left. White squares indicate regions displayed at higher magnification. For all confocal microscopy panels, orthogonal z-stack views and magnified insets are provided. Experiments were performed in biological triplicate (n = 3). Data are presented as mean ± SEM and were analyzed by two-way ANOVA followed by Bonferroni post-hoc test (**p < 0.01, ****p < 0.0001).

Although entotic structures in narciclasine-treated cells appeared smaller and more rounded, we did not detect clear differences in nuclear morphology or lysosomal accumulation when compared with control entotic structures. We therefore focused subsequent analyses on narciclasine-treated conditions.

To characterize distinct stages of entosis, we performed correlative light–electron microscopy (CLEM) in cells treated with 1 µM narciclasine and stained with Hoechst and Lysotracker red. Consistent with our light-microscopy analyses, narciclasine induced complex entotic structures that could be resolved ultrastructurally. CLEM enabled identification of multiple stages of entotic progression, including an early stage characterized by active invasion or potential exit of one cell within another, although precise directionality was difficult to determine (Fig. 3D). The observed ultrastructural morphology was highly consistent with entotic internalization, defined by rounded cellular architecture, the presence of a membrane-bound IC enclosed within an entotic vacuole, and the absence of apoptotic fragmentation or nuclear condensation, which would otherwise indicate apoptotic engulfment (Fig 3D). An intermediate stage was characterized by a cell internalized within an EC itself contained a large entotic vacuole, suggesting a nested entotic event and potential degradation of the internalized cell (Fig. 3E), as well as late-stage structures consistent with advanced digestion or near-complete clearance of the IC (Fig. 3F). Additional configurations included internal cell division, with two cells enclosed within a single EC (Supplementary Fig. 4A), and entosis in binucleated EC (Supplementary Fig. 4B). This observation is consistent with binucleation occurring downstream of entosis, although direct temporal evidence is not available in the current dataset.

Finally, in MCF7 cells expressing LC3–GFP, narciclasine treatment promoted formation of a ring-like LC3 structure at the entotic vacuole interface, consistent with previously described LC3-associated entotic vacuoles^30^ and supporting the conclusion that narciclasine induces bona fide entotic structures (Supplementary Figure 4C).

### Narciclasine induces entosis through ROCK1-mediated phosphorylation of MLC2

Next, we investigated whether ROCK1 activity is required for the effects of narciclasine on entotic formation. To this end, we combined narciclasine treatment with pharmacological inhibition of ROCK1 using the ROCK inhibitor Y27632, which is widely used to block entosis. As expected, treatment with Y27632 alone reduced basal entotic events compared with vehicle control (Fig. 4A and B). Consistent with our previous observations, narciclasine treatment increased entotic structure formation relative to vehicle. Notably, pre-treatment with Y27632 completely abolished narciclasine-induced CIC formation, indicating that ROCK activity is required for narciclasine-mediated induction of entosis (Fig. 4A and B).

**Figure 4.**
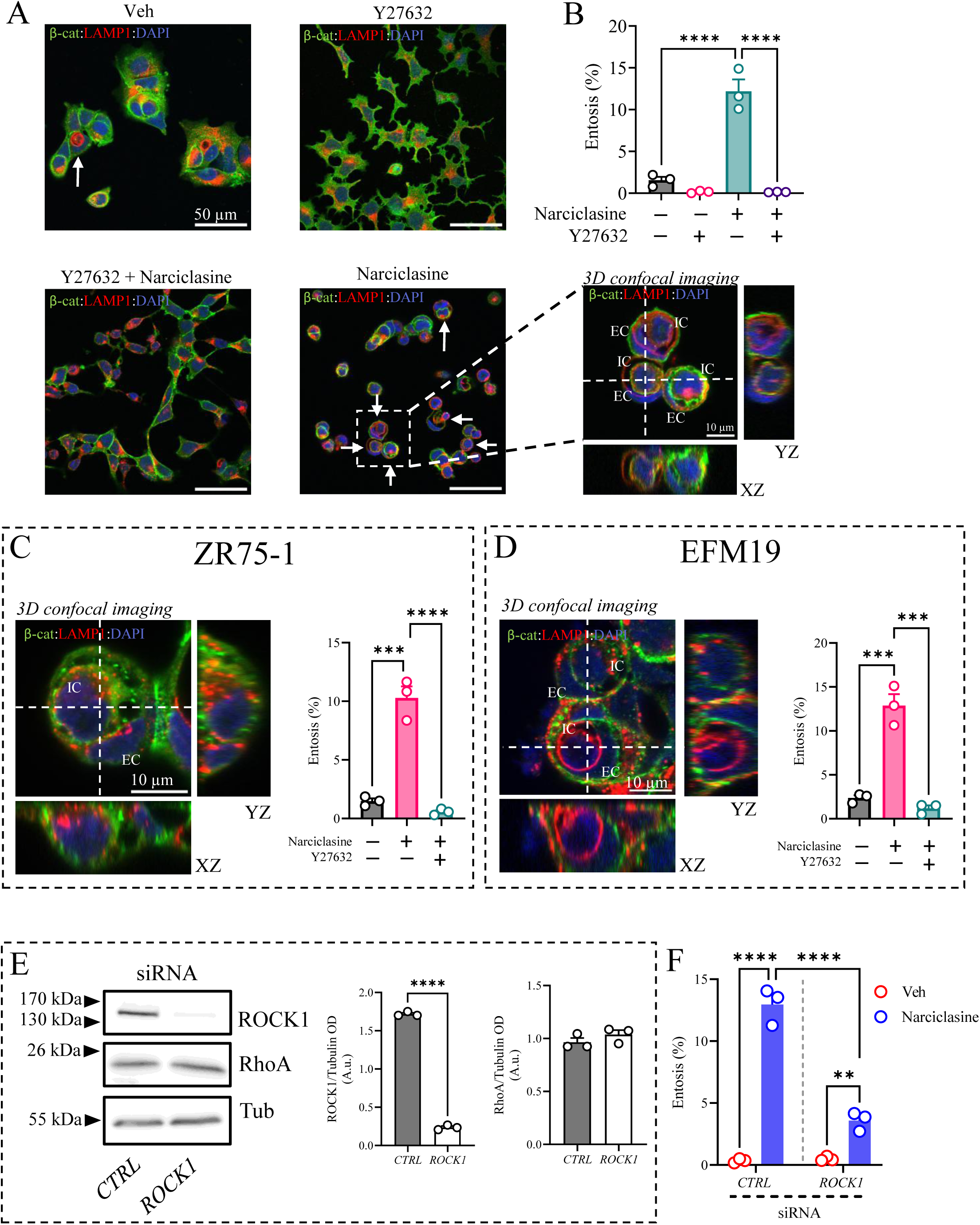
Pharmacological or genetic inhibition of ROCK1 abrogates narciclasine-induced entosis. **(A)** MCF7 cells were treated with vehicle (DMSO), 1 µM narciclasine, 20 µM Y27632, or pre-treated with 20 µM Y27632 followed by 1 µM narciclasine for 24 h. Entotic structures were visualized by immunofluorescence staining for β-catenin (membrane, green), LAMP1 (lysosomal compartments, red), and DAPI (nuclei, blue), followed by z-stack confocal microscopy. Representative 3D confocal images are shown. White arrows indicate entotic structures. Selected regions (boxed) are magnified, with corresponding orthogonal z-stack views displayed adjacent to each image. **(B)** Summary of quantification of entotic events in MCF7 cells treated from Figure A. **(C–D)** Equivalent analyses performed in ZR75-1 (C) and EFM19 (D) cells. Panels show representative entotic structures following treatment with 1 µM narciclasine and the corresponding quantification. **(E)** Representative immunoblot images of ROCK1, RhoA, and tubulin (loading control) in control or ROCK1-silenced cells, together with densitometric quantification (absolute OD unit) of ROCK1 and RhoA protein levels normalized to tubulin. **(F)** Quantification of entotic events in control and ROCK1-silenced MCF7 cells treated with vehicle or 1 µM narciclasine. For all confocal microscopy panels, orthogonal z-stack views and magnified insets are provided. Experiments were performed in biological triplicate (n = 3). Data are presented as mean ± SEM. Statistical analysis for (B–D) was performed using one-way ANOVA followed by Bonferroni post-hoc test (***p < 0.001, ****p < 0.0001), whereas panel E was analysed using a paired two-tailed Student’s t-test (****p < 0.0001). Panel F was analyzed using two-way ANOVA followed by Bonferroni post-hoc test (**p < 0.01, ****p < 0.0001).

We extended these findings to additional ER-positive breast cancer cell lines. In ZR-75-1 and EFM-19 cells, narciclasine similarly induced entotic structures formation, reaching approximately 10% and 13%, respectively (Supplementary Fig. 5 and 6; Fig. 4C and D). As in MCF7 cells, pre-treatment with Y27632 completely blocked narciclasine-induced entosis formation in both cell lines (Fig. 4C and D).

To further validate the requirement for ROCK1, we performed genetic silencing using a selective siRNA against ROCK1. ROCK1 knockdown led to a marked reduction in ROCK1 protein levels without affecting RhoA expression (Figure 4E). We then treated control and ROCK1-depleted cells with narciclasine and quantified CIC formation. In control siRNA conditions, narciclasine increased entotic events to approximately 15% (Figure 4F). In contrast, ROCK1 silencing significantly reduced narciclasine-entotic formation compared with control siRNA (Figure 4F), confirming that ROCK1 is required for narciclasine-mediated induction of entosis.

Following this, we sought to determine whether the RhoA–ROCK1 pathway was activated. To this end, we analyzed the accumulation of pMLC2, a direct ROCK1 substrate that is also involved in the early stages of entotic invasion. Time-dependent western blot analysis revealed an increase in pMLC2 upon treatment with 1 µM narciclasine (Fig. 5A and B), reaching a peak at 6 h followed by a reduction in signal at 16 h (Fig. 5A and B). No significant changes were detected in total RhoA or ROCK1 protein levels. However, the appearance of a lower-molecular-weight band below the ROCK1 signal suggests proteolytic cleavage associated with ROCK1 activation (Fig 5A). These findings were supported by immunofluorescence analysis, which showed pMLC2 accumulation at the cortex opposite to the site of invasion (Fig. 5C). To further assess the role of myosin II activity and the contribution of actomyosin contractility when using narciclasine, we used blebbistatin, a myosin II inhibitor^31^. Whereas narciclasine alone increased CIC formation, blebbistatin pre-treatment abolished narciclasine-induced entosis (Supplementary Fig. 7 and Fig. 5D), supporting a requirement for actomyosin contractility downstream of ROCK1 activation. Given the rapid induction of pMLC2 by narciclasine, we next examined whether this response coincided with early entotic onset following 6 h narciclasine treatment. Consistent with this possibility, treatment with 1 µM narciclasine increased the frequency of entotic structures to approximately 10% at this time point, when compared to vehicle (Fig. 5E and F).

**Figure 5.**
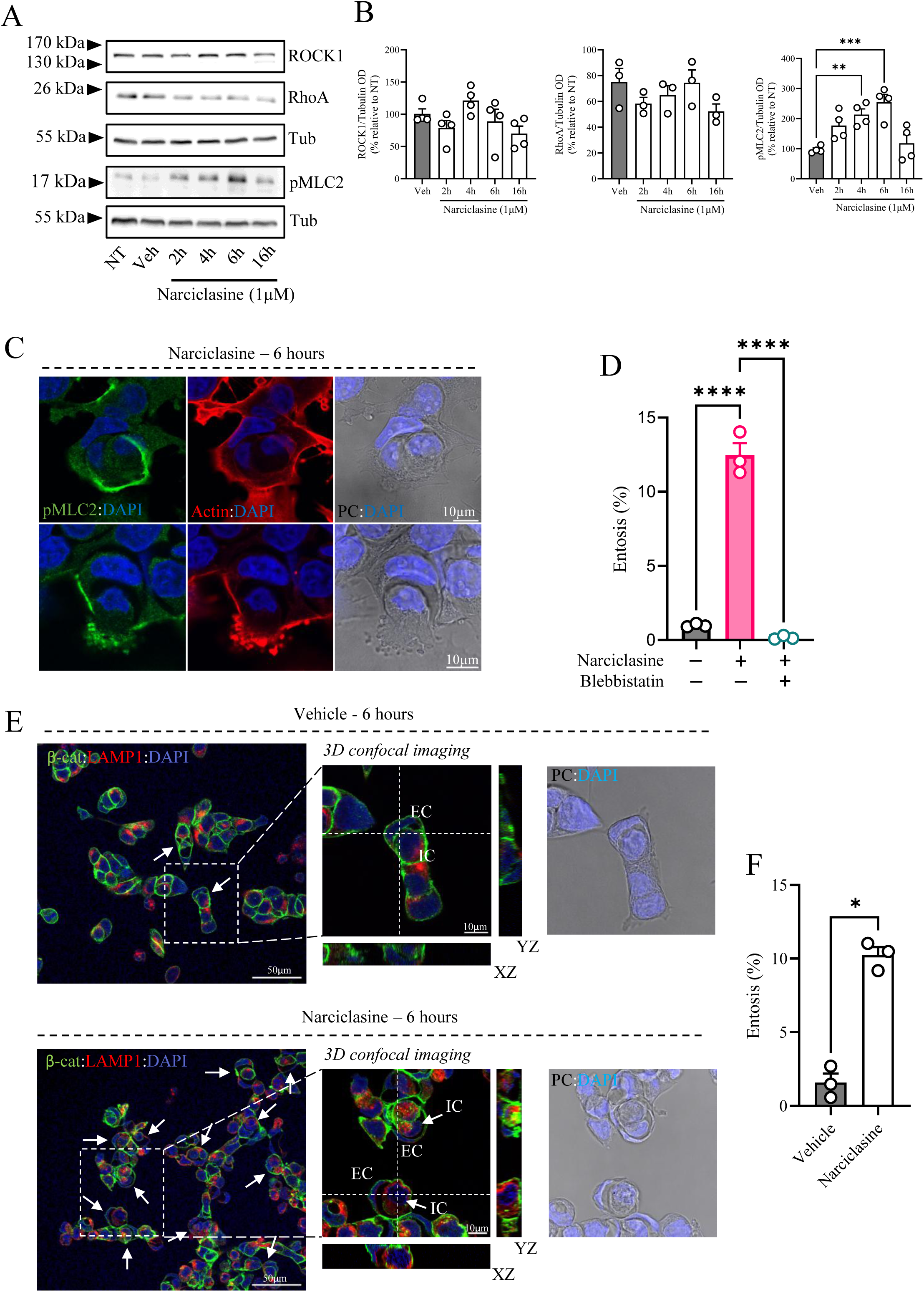
Narciclasine induces entosis within 6 hours through ROCK1-mediated phosphorylation of MLC2. **(A)** Representative immunoblot images of ROCK1, RhoA, pMCL2 and tubulin (loading control) in MCF7 treated with 1uM narciclasine at different time-points. **(B)** Densitometric quantification of ROCK1, RhoA and pMCL2 protein levels from immunoblot images. **(C)** Immunofluorescence staining for pMCL2 (green) and actin (red) in MCF7 cells treated with 1 µM narciclasine for 6 hours. Nuclei were counterstained with DAPI (blue). **(D)** Summary of quantification of entotic events in MCF7 cells treated with vehicle, 1 µM narciclasine, or 50 µM blebbistatin (1 hour pretreatment) in combination with 1 µM narciclasine for 24 hours. **(E)** Representative 3D confocal images of entotic structures following 6 hours of treatment with vehicle or 1 µM narciclasine. White arrows indicate entotic events. Selected regions (boxed) are magnified, with corresponding orthogonal z-stack views shown adjacent to each image. **(F)** Summary of quantification of entotic events from the conditions shown in panel E. For all confocal microscopy panels, orthogonal z-stack views and magnified insets are provided. Experiments were performed in biological triplicate (n = 3). Data are presented as mean ± SEM. Data in panel B are normalized to untreated controls within each experiment (set to 100%). Statistical analysis for panels B and D was performed using one-way ANOVA followed by Bonferroni’s post hoc test (**p < 0.01, ***p < 0.001, *p < 0.0001), whereas panel F was analyzed using a paired two-tailed Student’s t-test (*p < 0.05).

ROCK1 can be activated through caspase-3–mediated cleavage at the DETD 1113/G site, which removes its autoinhibitory domain and generates a constitutively active kinase^32^. We therefore investigated whether this proteolytic activation mechanism contributes to the initiation of entosis mediated by narciclasine. To this end, cells were pre-treated with the pan-caspase inhibitor zVAD-fmk prior to narciclasine exposure, and entotic events were quantified.

As previously observed, treatment with 1 µM narciclasine significantly increased entotic events formation (Supplementary Fig. 8; Figures 6A and B). Importantly, pre-treatment with zVAD-fmk did not significantly alter the frequency of entotic structures compared with narciclasine alone, with only a minor, non-significant increase observed (Figures 6B). These data indicate that caspase activity is not responsible for ROCK1 cleavage and activation upon narciclasine treatment and is not required for narciclasine-induced CIC formation.

**Figure 6.**
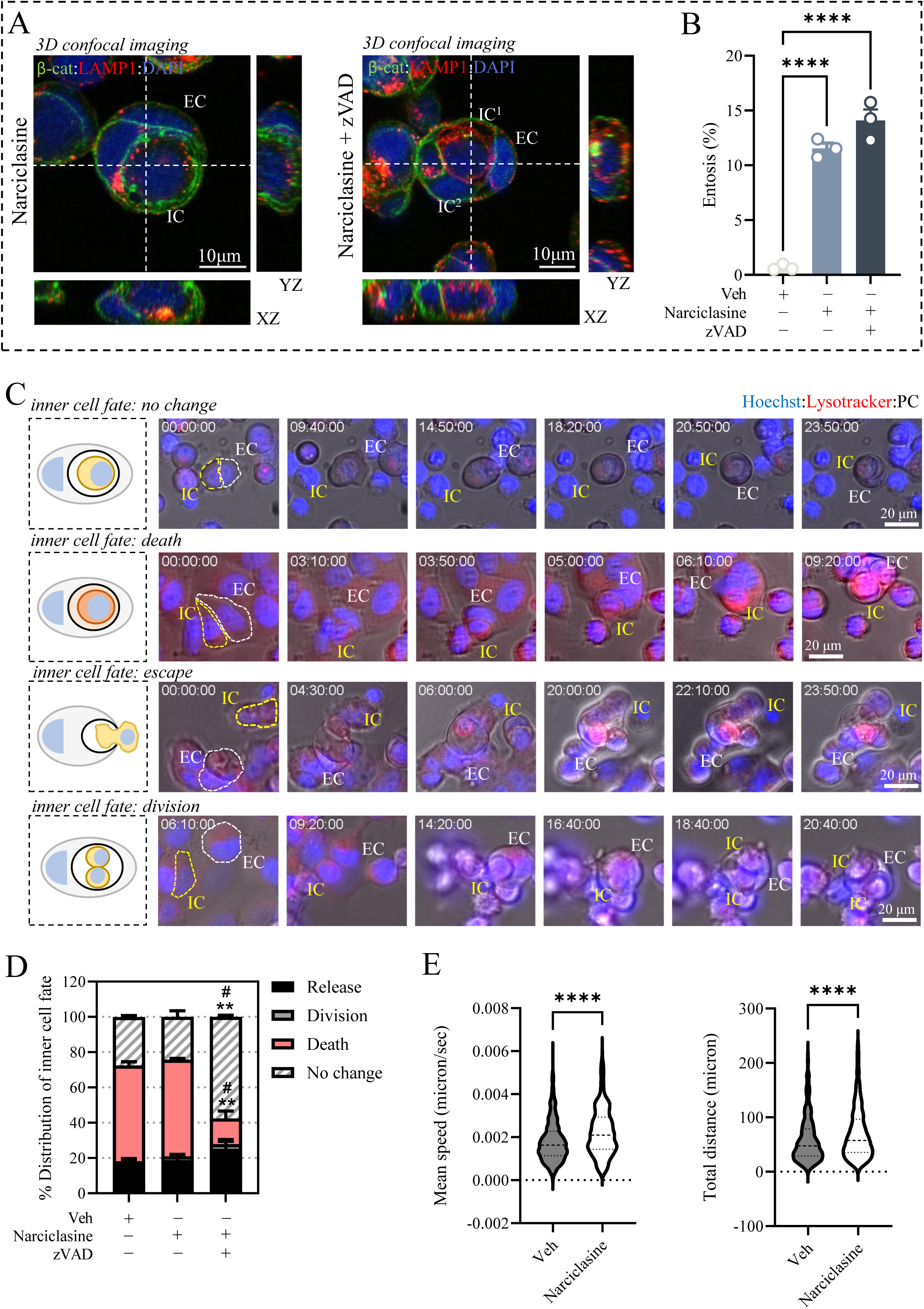
Caspase-mediated ROCK activation is not involved in narciclasine-induced entosis. **(A)** MCF7 cells were treated with vehicle (DMSO), 1 µM narciclasine, 50 µM zVAD, or pre-treated with 50 µM zVAD followed by 1 µM narciclasine for 24 h. Entotic structures were visualized by immunofluorescence staining for β-catenin (membrane, green), LAMP1 (lysosomal compartments, red), and DAPI (nuclei, blue), followed by z-stack confocal microscopy. Representative 3D confocal images are shown with corresponding orthogonal z-stack views displayed adjacent to each image. **(B)** Summary of quantification of entotic events in MCF7 cells treated from Figure A. **(C)** Time-lapse sequences of MCF7 cells stained with Lysotracker red (lysosomal compartments, red), and Hoechst (nuclei, blue) and treated with 1 µM narciclasine illustrating distinct inner cell (IC) fates, including persistence without change, entotic cell death (ECD), escape, and division. **(D)** Summary of quantification of IC fate distribution in MCF7 cells treated with vehicle, 1 µM narciclasine, or 50 µM zVAD in combination with 1 µM narciclasine. **(E)** Mean speed and total distance traveled, quantified from time-lapse imaging of MCF7 cells treated with vehicle or 1 µM narciclasine. Experiments were performed in biological triplicate (n = 3). Data are presented as mean ± SEM for panels B and D, and as median ± IQR for panel E. Statistical analysis for panel B was performed using one-way ANOVA followed by Bonferroni’s post hoc test (****p < 0.0001). Panel D was analyzed using two-way ANOVA followed by Bonferroni’s post hoc test (*p < 0.05, **p < 0.01); asterisks indicate significance between vehicle and the narciclasine + zVAD condition, whereas hash symbols (#) indicate significance between narciclasine alone and narciclasine + zVAD. Panel E was analyzed using an unpaired two-tailed Student’s t-test (****p < 0.0001).

To further characterize the fate of internalized cells under these conditions, we performed 24-hour time-lapse imaging and classified four distinct outcomes: no change (IC remains viable and quiescent), entotic cell death (ECD), division of the IC, and escape of the IC from the EC (Figure 6C). In vehicle-treated conditions, the majority of IC underwent ECD, followed by escape, with a smaller fraction displaying no change, and only a very minor proportion undergoing division (Figure 6D). Similar distributions of IC fates were observed following narciclasine treatment (Figure 6D). Notably, pre-treatment with zVAD-fmk markedly altered the fate of IC. Under these conditions, the majority of IC exhibited no change, followed by escape, while only a small fraction underwent ECD (Figure 6D).

In addition, based on qualitative observations of altered cellular dynamics, we quantified cell motility parameters. Narciclasine-treated cells exhibited a significant increase in both migration speed and total distance traveled compared with vehicle-treated controls (Figure 6E), indicating that narciclasine enhances cellular motility, which may contribute to entosis.

### Pharmacological ROCK1 activation reprograms the proteome toward cytoskeletal remodeling and adhesion machinery, while suppressing proliferative and metabolic regulators

To better understand the molecular consequences of narciclasine treatment, we performed bulk proteomic analysis followed by differential expression profiling in MCF7 cells treated with vehicle or 1 µM narciclasine for 24 hours. Quantitative proteomic profiling of narciclasine-treated cells revealed selective and functionally coherent remodelling rather than global proteotoxic disruption, as visualized by volcano plot analysis of differential protein abundance (Fig. 7A). Among the most significantly up-regulated proteins (Supplementary Table 1 and Fig. 7A) were the Ca²⁺-dependent membrane scaffold Annexin-A6 (ANXA6), the actin-bundling factors PLS2 (also called LCP1) and Plastin-3 (PLS3, also termed T-Plastin), and the extracellular matrix cross-linking enzyme lysyl oxidase (LOX), collectively defining a mechanics- and adhesion-associated molecular signature consistent with enhanced ROCK1-dependent cytoskeletal tension. Additional increases in vesicular trafficking and membrane organization regulators, including VPS54, LZTFL1, and phosphoinositide phosphatases PIP4P1/2, further support activation of membrane remodeling programs required for entotic internalization. We also observed elevation of metabolic and redox-associated enzymes such as MLYCD, PYROXD2, MAOB, DDAH1/2, and SELENBP1 that indicates metabolic adaptation rather than nonspecific cytotoxic stress. In contrast, the most strongly down-regulated proteins (Suppementary Table 2 and Fig. 7A) were enriched for regulators of cell-cycle progression, mitochondrial metabolism, and lineage identity, including Cyclin D1 (CCND1), CDK4, the cholesterol biosynthesis enzyme HMGCR, mitochondrial factors (COX16, DNAJC15, CHCHD2), and the luminal breast lineage transcription factor GATA3. Suppression of proliferative drivers together with attenuation of mitochondrial and biosynthetic enzymes indicates a shift away from anabolic growth toward a mechanically engaged cellular state. Notably, reduction of the microtubule-associated organizer MTCL2 and additional chromatin-associated regulators further supports broad cytoskeletal and structural reprogramming.

**Figure 7.**
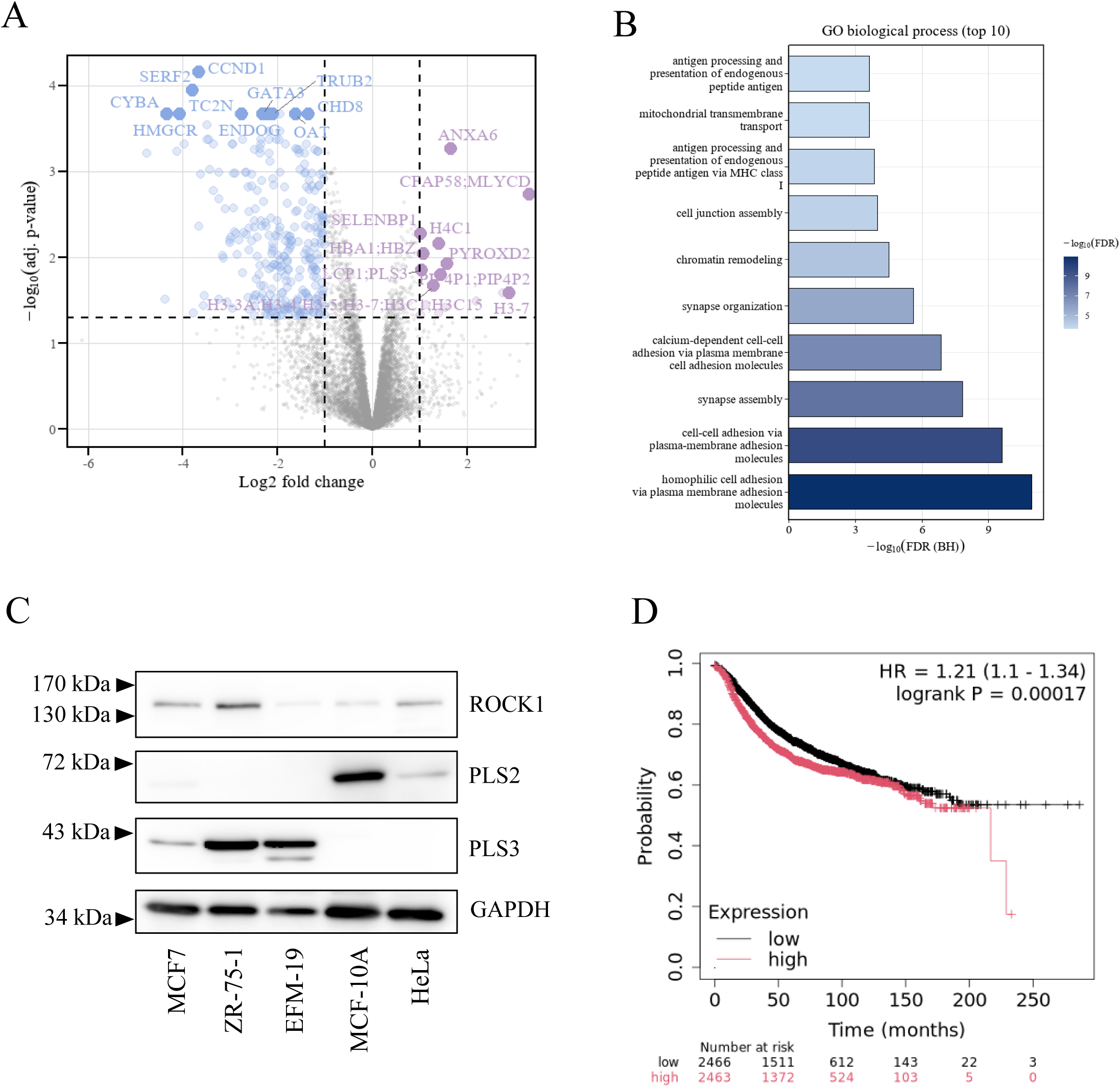
Proteomics analysis of MCF7 treated with narciclasine reveals Cytoskeletal regulators. **(A)** Volcano plot of proteomic changes following narciclasine treatment, displaying log₂ fold change versus −log₁₀ adjusted *P* value. The 10 most significantly up- and downregulated proteins are highlighted and annotated. **(B)** Gene Ontology (GO) biological process enrichment analysis of differentially expressed proteins. The top 10 enriched terms are shown, ranked by −log₁₀(FDR), with enrichment prominently observed in cell–cell adhesion, plasma membrane adhesion, and homophilic adhesion pathways. **(C)** Immunoblot analysis of ROCK1, PLS2, and PLS3 protein expression across epithelial and breast cancer cell lines (MCF7, ZR-75-1, EFM-19, MCF-10A, and HeLa), with GAPDH as loading control. **(D)** Kaplan–Meier survival analysis stratified by PLS3 expression, showing decreased survival probability in patients with high PLS3 expression compared with low expression. Numbers at risk are indicated below the plot.

Functional interpretation of significantly altered proteins using Gene Ontology Biological Process enrichment demonstrated dominant over-representation of cell–cell adhesion, plasma-membrane adhesion molecule interactions, synapse assembly, and calcium-dependent adhesion pathways, identifying adhesion remodeling as the principal biological output of narciclasine exposure (Fig. 7B). Complementary preranked Reactome enrichment analysis independently revealed coordinated activation of RHO GTPase signaling, ROCK-dependent cytoskeletal regulation, integrin signaling, actin assembly, and junctional trafficking pathways, defining a coherent mechanical signaling network centered on actomyosin contractility (Supplementary Fig. 9A).

### PLS3 expression identifies breast cancer cells susceptible to narciclasine-induced entosis and associates with poorer clinical outcomes

Among the proteins identified as significantly up- or down-regulated, we prioritized candidates involved in actin cytoskeleton dynamics, given their potential relevance to the mechanical processes underlying entosis. In particular, we focused on PLS3 and PLS2, two calcium dependent actin-bundling proteins known to regulate filament organization, cortical tension, and cell motility. We first examined the basal expression levels of these proteins across the panel of cell lines used for narciclasine treatment. While ROCK1 expression varied between cell lines, a distinct pattern emerged for the actin regulators. The breast cancer cell lines MCF7, ZR75-1, and EFM19 exhibited clear accumulation of PLS3, whereas the non-transformed epithelial line MCF10A showed low to undetectable PLS3 levels. Conversely, PLS2 was low or absent in the breast cancer cell lines but displayed higher abundance in MCF10A (Fig. 7C). Based on this differential expression profile, we next evaluated the effect of narciclasine in MCF10A cells. As shown in Supplementary Figure 9B and C, narciclasine treatment did not increase entosis compared with vehicle control, in contrast to the response observed in breast cancer cells highlighted in Fig 4.

To evaluate the potential clinical relevance of PLS3 in breast cancer -without aiming to define a biomarker- we interrogated publicly available transcriptomic datasets^33^ comprising pooled microarray cohorts. Patients were stratified according to median PLS3 expression, and relapse-free survival (RFS) was assessed across all molecular subtypes without additional restrictions. Interestingly, high PLS3 expression was associated with significantly reduced RFS compared to low-expression tumors (hazard ratio [HR] = 1.21, 95% CI 1.10–1.34; log-rank P = 0.0002). The separation between survival curves was evident early and remained consistent over long-term follow-up (Fig. 7D). Although the effect size was moderate, the robustness of the association across a large pooled cohort suggests that elevated PLS3 expression correlates with increased risk of disease recurrence in breast cancer.

### Narciclasine causes recruitment of PLS3 to the plasma membrane, which triggers entosis

Based on the differential expression of PLS3 observed in our proteomic analysis, we next sought to validate these findings and assess their functional relevance. Western blot analysis confirmed that narciclasine treatment increased PLS3 protein levels after 24 hours (Supplementary Fig. 10A). In parallel, immunofluorescence analysis revealed a marked redistribution of PLS3, shifting from a diffuse cytoplasmic pattern under vehicle conditions to a membrane-enriched localization following narciclasine exposure (Fig. 8A and B). Figure 8A also shows that PLS3 localizes to the plasma membrane only in regions lacking cell–cell contacts.

**Figure 8.**
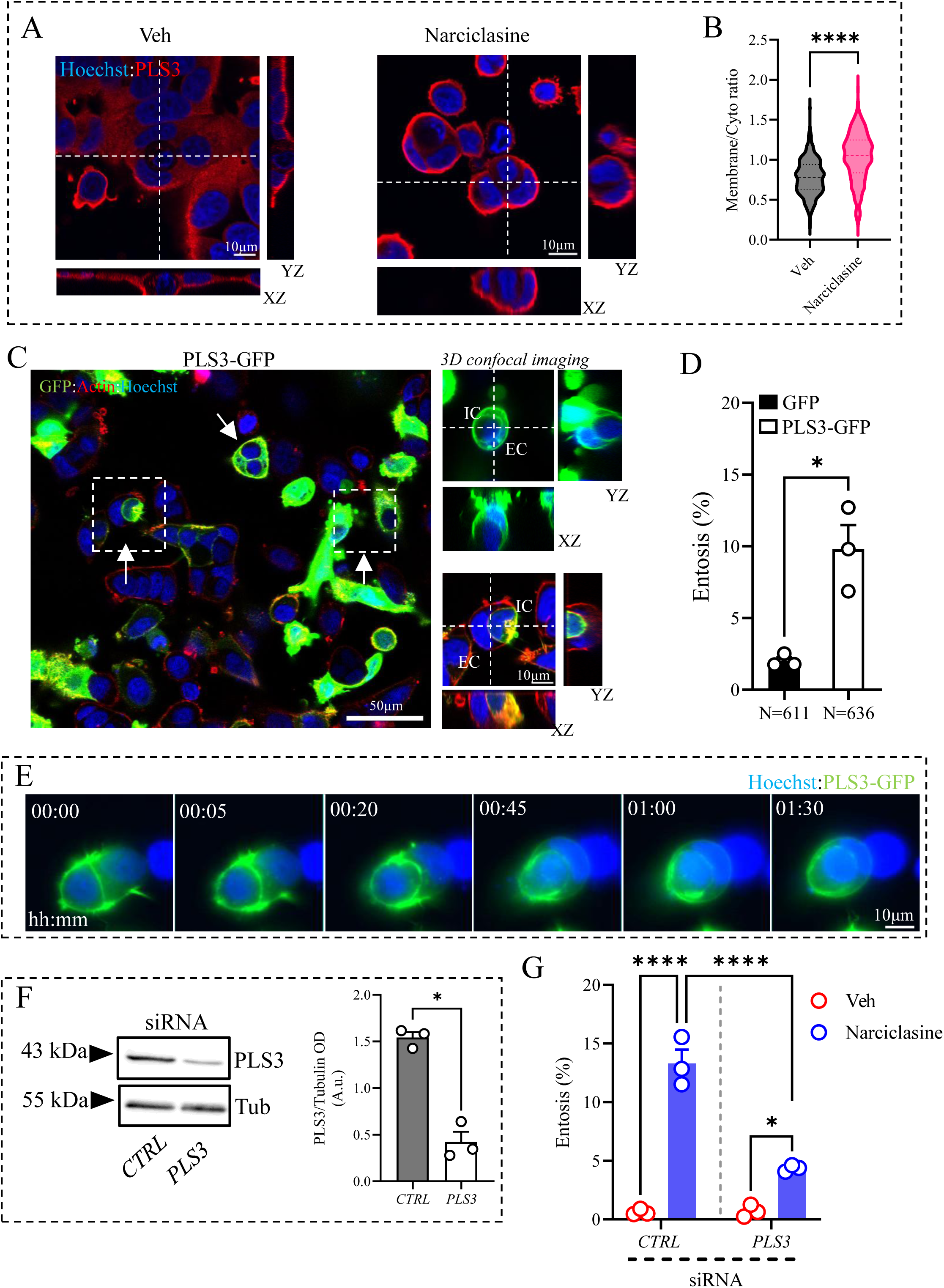
PLS3 is recruited to the plasma membrane to trigger entosis. **(A)** Immunofluorescence staining for PLS3 (red) in MCF7 cells treated with 1 µM narciclasine or vehicle for 24 hours. Nuclei were counterstained with DAPI (blue). **(B)** Summary of quantification of membrane to cytosol ratio (mean intensity) from immunofluorescence PLS3 images. **(C)** Representative 3D confocal images of live MCF7 cells expressing PLS3-GFP and stained with SiR-Actin (red) and Hoechst (blue). White arrows indicate entotic structures. Boxed regions showing representative entotic structures are magnified, with corresponding orthogonal z-stack views shown alongside each image. **(D)** Quantification of entotic events in MCF7 cells expressing GFP or PLS3-GFP. **(E)** Representative time-lapse images (GFP and Hoechst) showing induction of entosis in MCF7 cells expressing PLS3. **(F)** Representative immunoblot images of PLS3 and tubulin (loading control) in control or PLS3-silenced cells, together with densitometric quantification of PLS3 normalized to tubulin. **(G)** Summary of quantification of entotic events in control and *PLS3*-silenced MCF7 cells treated with vehicle or 1 µM narciclasine. For all confocal microscopy panels, orthogonal z-stack views and magnified insets are provided. Experiments were performed in biological triplicate (n = 3). Data are presented as median ± IQR for panel B and as mean ± SEM for panels D, F and G. Statistical analysis for panel B was performed using an unpaired two-tailed Student’s t-test (****p < 0.0001), whereas panel D and F by a paired two-tailed Student’s t-test (*p < 0.05). Panel G was assessed by a two-way ANOVA followed by Bonferroni’s post hoc test (*p < 0.05, ****p < 0.0001).

To determine whether PLS3 contributes to entotic structure formation, we overexpressed a PLS3-GFP construct in MCF7 cells. Overexpressed PLS3 accumulated prominently at the cell membrane, and quantification restricted to GFP-positive cells demonstrated a significant increase in entotic structures compared with control GFP-transfected cells (Supplementary Figure 10B and C; Fig. 8C and D). Time-lapse imaging further confirmed active cellular invasion events consistent with entosis (Fig. 8E, Supplementary Figure 10D and Video 7).

Conversely, when PLS3 was silenced (Fig. 8F) we observed a markedly impaired narciclasine-induced entosis (Fig 8G). While narciclasine robustly increased entotic frequency in control siRNA conditions, PLS3 depletion reduced entosis to ∼4%, indicating that PLS3 is required for the full entosis-inducing effect of narciclasine (Fig. 8G).

## DISCUSSION

Entosis represents a driven form of CIC invasion governed by E-cadherin–based adherens junctions and actomyosin contraction^34,35^, primarily controlled by RhoA–ROCK signaling and related GTPases. Two key challenges continue to limit mechanistic understanding of entosis. First, despite nearly two decades of research, the molecular determinants linking actin remodeling and membrane dynamics during entotic invasion remain incompletely defined, and the cytoskeletal mechanisms that physically enable one cell to invade another are still not fully understood. Second, progress in the field has been hindered by the lack of experimental strategies to robustly induce entosis without simultaneously introducing cellular stresses that may activate additional signaling pathways, complicating precise mechanistic interrogation of the internalization process. In this context, identifying conditions that reproducibly trigger the entotic internalization program while allowing mechanistic interrogation of the underlying cytoskeletal machinery remains an important challenge. In this study, we identify a pharmacologically targetable ROCK1 axis that is sufficient to induce entosis in breast cancer cells and define PLS3 as a critical downstream effector enabling entosis initiation.

We first demonstrate that constitutive activation of ROCK1 is sufficient to trigger entotic invasion, as expression of a constitutively active ROCK1 construct increased entosis frequency and recapitulated the canonical morphological sequence of entosis observed by live-cell imaging. These findings establish enforced ROCK1 signaling as a driver of the mechanical program required for cell internalization, consistent with prior work positioning RhoA–ROCK–myosin contractility at the core of entotic invasion^16,34^. This observation aligns with previous reports showing localized RhoA activation at the distal cortex of internalizing cells and the requirement for RhoA–ROCK activity during invasion and also with the notion that ROCK kinases are required only for early-middle stage progression^34,35^. Notably, this differs slightly from earlier studies reporting that ROCK1 overexpression alone was sufficient to confer loser status in mixed populations^36^, as GFP-ROCK1 –expressing cells were detected in both internal and external positions. In our hands, these findings instead correlate with the requirement for ROCK1 activation rather than simple increases in ROCK1 abundance. Although RhoA activation is typically spatially restricted to the cortex opposite the site of invasion, our data indicate that global ROCK1 activation alone is sufficient to significantly increase entotic structures. More precise spatiotemporal activation of RhoA or ROCK1 will be required to determine whether selective mechanical polarization can further refine or enhance entosis induction.

We next show that narciclasine induces entosis formation, dynamic invasion behaviors, and ultrastructurally defined entotic architectures across a range of concentrations. Of particular interest is the emergence of complex entotic architectures, including nested configurations containing three cells or CIC structure inside a single cell. This may reflect strong activation of the ROCK1 pathway across the population, enforcing repeated or sequential internalization events and generating diverse CIC morphologies.

Mechanistically, multiple lines of evidence demonstrate that ROCK1 activity is required for narciclasine-induced entosis. Pharmacological inhibition with Y27632 abolished CIC formation across several ER-positive breast cancer cell lines, while genetic silencing of ROCK1 partially reduced this effect. The incomplete suppression may reflect compensatory activity from ROCK2, which was not directly examined in this study. Consistent with ROCK pathway engagement, narciclasine rapidly increased phosphorylation of MLC2—a direct ROCK substrate linked to cortical contractility and early entotic invasion^36^—and inhibition of myosin II activity prevented CIC formation, consistent with previous reports^16^. Together, these findings position actomyosin contractility downstream of narciclasine and upstream of entotic internalization, defining a coherent mechanical signaling cascade.

Previous studies reported that narciclasine activates Rho signaling and promotes stress-fiber formation in glioblastoma and endothelial cells^28,37^, in some contexts through partially RhoA-independent ROCK activation^37^. In our system, narciclasine induced a peak in pMLC2 levels at approximately 6 h, coinciding with the early appearance of entosis (∼10%), indicating that the mechanical response to this compound is rapid. The decline in pMLC2 observed at 16 h is consistent with a model in which ROCK-dependent myosin activation functions as an early initiating event rather than a persistent late-state marker. Given the asynchronous nature of entosis in cell populations, bulk biochemical measurements at later time points likely underestimate localized or transient contractile signaling associated with the initial invasion step.

While approaches enabling acute and spatially restricted ROCK1 activation will further refine this model, the rapid ROCK-dependent induction of entosis by pharmacological stimulation—together with suppression by ROCK inhibition or ROCK1 depletion—demonstrates the functional sufficiency of ROCK pathway activation to initiate entotic CIC invasion. It is worth noting that although narciclasine robustly induced ROCK1 activation in our system, whether this effect occurs through direct engagement of ROCK1 or indirectly via upstream signaling pathways remains to be determined.

Notably, narciclasine-induced entosis formation occurred independently of caspase activation, excluding indirect contributions from apoptosis-associated detachment or secondary engulfment. Time-lapse imaging instead revealed canonical entotic fates, including death, escape, quiescence, and division. We observed differences in Lysotracker accumulation between Fig. 3A,B and Fig. 6D. In fixed time-point experiments, detection of ECD depends on capturing cells at the appropriate stage of entosis, which is less likely in vehicle-treated conditions where entosis occurs at low frequency and progresses asynchronously. In contrast, narciclasine increases the number of entotic events and likely synchronizes their progression, making Lysotracker-positive ECD cells easier to detect.

Additionally, we also observed increased cellular motility following narciclasine exposure. This observation is particularly relevant because increased cellular motility is closely associated with invasive behavior^38^ and is consistent with enhanced ROCK1 signaling. Indeed, NRF2-dependent upregulation of the RhoA–ROCK1 axis has been shown to promote cell motility^39^. Additionally, activating mutations in ROCK1 that elevate kinase activity drive actin cytoskeletal remodeling, resulting in increased motility accompanied by reduced cell–cell adhesion^40^. Following on, combined treatment with zVAD and narciclasine increased the proportion of cells not undergoing ECD. Although entotic death is considered caspase-independent, similar effects have been reported previously, suggesting indirect crosstalk between apoptotic inhibition and entotic fate decisions^19^.

Global proteomic profiling further supports the above mentioned results by revealing selective remodeling of cytoskeletal, adhesion, and membrane-trafficking networks, accompanied by suppression of proliferative and metabolic regulators. Enrichment of RHO GTPase signaling, ROCK-dependent cytoskeletal regulation, integrin signaling, and adhesion pathways indicates that narciclasine induces a mechanically engaged cellular state rather than generalized toxicity. Within this program, the actin-bundling proteins PLS3 and PLS2 emerged as candidate mediators linking ROCK1 activation to cytoskeletal reorganization. Profiling across cancer cell lines revealed higher PLS3 accumulation in breast tumor cells relative to PLS2, whereas non-transformed mammary epithelial cells displayed the opposite pattern and remained unresponsive to narciclasine. These findings suggest that tumor-specific mechanical priming may be required for pharmacological induction of entosis.

PLS3 (T-plastin) is broadly expressed in solid tissues^41^ and has been repeatedly associated with epithelial tumor biology^42,43^. In breast cancer, elevated PLS3 expression has been associated with increased metastatic potential and has been proposed as a marker of circulating tumor cells (CTCs) in epithelial malignancies. Importantly, patients with PLS3-positive tumors exhibit significantly reduced overall survival and disease-free survival compared with PLS3-negative patients^44^. In contrast, PLS2 (LCP1 or L-plastin) is predominantly expressed in hematopoietic and immune cells under physiological conditions and plays established roles in leukocyte activation, adhesion, and immune synapse formation^45^. Although ectopic PLS2 expression has been reported in some solid tumors^46,47^, its canonical biological context remains immune-restricted, suggesting that its functional contribution in epithelial cancer may be secondary or context-dependent.

In addition to the main candidates, the proteomic analysis revealed several other proteins of potential relevance: LZTFL1, originally described as a regulator of ciliary trafficking and BBSome function^48^, has more recently been implicated in epithelial integrity and E-cadherin stabilization at the plasma membrane^49^. Annexin A6 (ANXA6) adds an additional layer of regulation at the membrane–cytoskeleton interface. As a Ca²⁺-dependent phospholipid-binding protein, Annexin A6 modulates membrane organization, cholesterol distribution, and cortical actin dynamics, and has been linked to E-cadherin trafficking and junctional stability^50–52^. Because entosis involves extensive membrane remodeling, vacuole formation, and dynamic reorganization of cortical tension, LZTFL1 and Annexin A6 could influence CIC formation by regulating membrane rigidity and actomyosin coupling at sites of cell–cell contact.

Given the above, we focused on PLS3; narciclasine increased PLS3 abundance and promoted its redistribution to the cell cortex, consistent with a role in actin bundling^53^. Functional studies demonstrate that PLS3 is required for narciclasine-induced entosis and is sufficient to enhance entotic formation upon overexpression, establishing PLS3 as a key cytoskeletal effector of the entotic program. It has been showed that PLS3 promotes cytoskeletal remodeling by cross-linking actin filaments into parallel and antiparallel bundles^54^, thereby stabilizing membrane protrusions and reinforcing lamellipodia and filopodia to support directional cell migration, particularly under mechanically challenging conditions such as matrix discontinuities^55^. Consistently, PLS3 is required for efficient lamellipodial protrusion and is enriched at the leading edge of migrating epithelial cells^56^, where it accelerates protrusion dynamics. Loss of PLS3 markedly reduces cell velocity and impairs wound closure in scratch assays, underscoring its functional role in migration and force transmission during invasion^55^. This actin-driven membrane protrusions that mediate invasive cell behaviors in other contexts could also be required for one cell to physically invade another.

Notably, PLS3 is a Ca²⁺-regulated actin-bundling protein, linking its activity to calcium-dependent control of actomyosin contractility. Because entosis critically depends on intracellular Ca²⁺ signaling—mediated in part by the SEPTIN–Orai1–Ca²⁺/CaM–MLCK axis that drives actomyosin-dependent engulfment^57^—these observations position PLS3 as a plausible mechanistic interface between calcium signaling, cytoskeletal remodeling, and entotic internalization. Together, these data position PLS3 downstream of ROCK1-mediated contractility and identify actin-bundling capacity as a limiting determinant of entotic internalization.

Previous work has shown that entotic invasion involves sustained plasma membrane blebbing controlled by the MRTF–SRF transcriptional pathway and its downstream effector Ezrin, linking actin dynamics to membrane remodeling during CIC invasion^58^. Conceptually, this study defines a pharmacologically inducible mechanical switch for entosis, linking ROCK1 activation to cytoskeletal remodeling and CIC formation through PLS3. These findings extend current models of entosis by demonstrating that chemical activation of contractile signaling is sufficient to initiate the process, and by identifying a specific actin-bundling effector required for efficient internalization. More broadly, the data support a model in which tumor cells exploit cytoskeletal plasticity to engage entotic behaviors that may influence cellular competition, survival, and therapeutic response within the tumor microenvironment.

These findings also open several avenues for future investigation. First, our analyses were primarily performed in in vitro breast cancer models, and it will be important to determine whether the ROCK1–PLS3 axis regulates entosis across additional tumor types and in physiological in vivo contexts. Second, although PLS3 is clearly required for narciclasine-induced CIC formation, the molecular mechanisms by which PLS3 coordinates cortical tension, adhesion remodeling, and membrane dynamics remain unresolved.

In summary, we identify ROCK1 activation as a sufficient and druggable trigger of entosis and uncover PLS3 as a critical cytoskeletal effector enabling CIC invasion. Together, these findings establish a mechanochemical pathway controlling entotic behavior in breast cancer cells and provide a conceptual framework for therapeutically modulating mechanically driven cell competition.

## Acknowledgments

The proteomic analysis was performed in the proteomics facility of SCSIE University of Valencia, a member of Proteored. We acknowledge the Unidad Mixta de Microscopía Electrónica CIPF-ISABIAL for providing access to electron microscopy facilities and technical support. We particularly thank Andrea Ibáñez Grau for her assistance with sample preparation, imaging, and expert guidance.

## Contributions

F.L and E.B. conceived the project. F.L. designed the experiments. F.L., S.P., C.G., A.H.C. and M.S. performed the experiments. F.L., S.P. and C.G. analysed all processed data. F.L. wrote the paper. E.B., S.P., L.A.N. J.A.G.S. and M.O.C. contributed to editing and revising the manuscript. All authors discussed the results and reviewed the paper.

## Supplementary Information

**Supplementary Figure 1. CIC structures in GFP-ROCK1 and GFP-ROCK1 Δ3 expressing cells.**

**(A)** Schematic representation of human ROCK1 and ROCK1 Δ3 protein. **(B)** Representative immunoblots showing GFP and tubulin (loading control) in MCF7 cells transfected with GFP, GFP–ROCK1, or constitutively active GFP–ROCK1 Δ3 after 24 h transfection. **(C)** Representative 3D confocal images of entotic structures in live MCF7 cells expressing GFP–ROCK1 and stained with SiR-Actin (red) and Hoechst (blue). **(D)** Representative 3D confocal images of entotic structures in live MCF7 cells expressing GFP–ROCK1 Δ3 and stained with SiR-Actin (red) and Hoechst (blue). Experiments were performed in biological triplicate (n = 3). Corresponding orthogonal z-stack views are shown alongside each image.

**Supplementary Figure 2. Narciclasine induces different types of entotic structures.**

**(A)** Representative time-lapse images (phase contrast, Hoechst and Lysotracker red) showing induction of entosis in MCF7 cells treated with 1 µM narciclasine. **(B)** Representative time-lapse images (phase contrast, Hoechst and Lysotracker red) showing an IC being released from an EC in MCF7 cells treated with 1 µM narciclasine. (**C)** Representative phase contrast, Hoechst and Lysotracker red) showing an IC undergoing ECD in MCF7 cells treated with 1 µM narciclasine. In all panels white arrows indicate IC and EC. **(D)** Representative phase contrast (PC) and DAPI confocal image for Figure 2D showing a one-cell-inside-another entotic structure in MCF7 cells treated with 1 µM narciclasine. **(E)** Representative phase contrast (PC) and DAPI confocal image for Figure 2E showing a nested entotic configuration (cell-in-cell-in-cell) in MCF7 cells treated with 1 µM narciclasine. **(F)** Representative phase contrast (PC) and DAPI confocal image for Figure 2F showing three cells internalized within a single EC following treatment with 1 µM narciclasine.

**Supplementary Figure 3. VEM analysis of CIC structures.**

**(A)** Representative transmission electron micrograph for vEM experiments showing multiple CIC structures following treatment of MCF7 with 1 μM narciclasine for 24 hours. Arrows indicate representative examples of ICs fully enclosed within EC. Inner cells display varying ultrastructural features consistent with different stages of entotic progression, including preserved nuclear morphology in early stages and cytoplasmic vacuolization and electron-dense degradation in later stages. ECs exhibit crescent-shaped nuclei and cytoplasmic remodeling around the ICs. **(B)** Volumetric electron microscopy image showing an EC with an enlarged entotic vacuole with a digested IC inside following treatment with 1 µM narciclasine. **(C)** Volumetric electron microscopy image of a classical CIC entotic structure in MCF7 cells treated with 1 µM narciclasine. For all electron microscopy panels, orthogonal z-stack views and magnified insets are provided. Experiments were performed in biological triplicate (n = 3).

**Supplementary Figure 4. Characterization of entotic structures in MCF7 cells treated with narciclasine.**

**(A)** CLEM analysis of two ICs inside an EC in MCF7 cells treated with 1 µM narciclasine (24 h). **(B)** CLEM analysis of two ICs inside an EC with failed kitokinesis in MCF7 cells treated with 1 µM narciclasine (24 h). In all panels are shown representative confocal images of Hoechst (blue) and LysoTracker (red) on the right; corresponding transmission electron microscopy (TEM) images are shown in the center, and merged correlative overlays are shown on the left. White squares indicate regions displayed at higher magnification. **(C)** Live-cell 3D confocal images of MCF7-LC3 GFP cells (stained with Hoechst) showing LC3 accumulation forming a ring-like structure around the entotic vacuole following treatment with 1 µM narciclasine for 24 h. For all confocal microscopy panels, orthogonal z-stack views and magnified insets are provided. Experiments were performed in biological triplicate (n = 3).

**Supplementary Figure 5. Narciclasine induces entosis in ZR75-1 cells.**

**(A)** Representative phase contrast (PC) and DAPI confocal image for Figure 4C. **(B)** Representative 3D confocal images of vehicle, narciclasine (1 µM) and narciclasine (1 µM) + Y27632 (20 µM, 1 hour pre-treatment) ZR75-1 treated cells (24 h), for figure 4C. Entotic structures were visualized by immunofluorescence staining for β-catenin (membrane, green), LAMP1 (lysosomal structures, red) and DAPI (nuclei, blue) followed by z-stack confocal microscopy. White arrows indicate entotic structures. Selected region (boxed) shows the representative entotic structure in Figure 4C. Shown are different Z positions.

**Supplementary Figure 6. Narciclasine induces entosis in EFM19 cells.**

**(A)** Representative phase contrast (PC) and DAPI confocal image for Figure 4D. **(B)** Representative confocal images of vehicle, narciclasine (1 µM) and narciclasine (1 µM) + Y27632 (20 µM, 1 hour pre-treatment) EFM19 treated cells (24 h), for figure 4D. Entotic structures were visualized by immunofluorescence staining for β-catenin (membrane, green), LAMP1 (lysosomal structures, red) and DAPI (nuclei, blue) followed by z-stack confocal microscopy. White arrows indicate entotic structures. Selected region (boxed) shows the representative entotic structure in Figure 4D. Shown are different Z positions.

**Supplementary Figure 7. Blebbistatin abolishes narciclasine entosis inducing effect.** Representative 3D confocal images of entotic events in MCF7 cells treated with vehicle, 1 µM narciclasine, or 50 µM blebbistatin (1 hour pretreatment) in combination with 1 µM narciclasine for 24 hours for quantification in Figure 5D. Entotic structures were visualized by immunofluorescence staining for β-catenin (membrane, green), LAMP1 (lysosomal structures, red) and DAPI (nuclei, blue) followed by z-stack confocal microscopy. White arrows indicate entotic structures. Selected region (boxed) shows a magnified representative FOV.

**Supplementary Figure 8. zVAD does not alter Narciclasine-induced entosis.**

**(A)** Representative phase contrast (PC) and DAPI confocal image for Figure 6A. **(B)** Representative 3D confocal images of vehicle, narciclasine (1 µM) and narciclasine (1 µM) + zVAD (50 µM pretreatment for 1 hour) MCF7 treated cells (24 h). Entotic structures were visualized by immunofluorescence staining for β-catenin (membrane, green), LAMP1 (lysosomal structures, red) and DAPI (nuclei, blue) followed by z-stack confocal microscopy. White arrows indicate entotic structures. Selected region (boxed) shows the magnified representative FOV in Figure 6A.

**Supplementary Figure 9. Narciclasine induces enrichment of cytoskeleton and cellular mechanics pathways in MCF7 cells.**

**(A)** Reactome pathway enrichment analysis (fgsea) of differentially expressed proteins identified by proteomics in MCF7 cells treated with 1 µM narciclasine for 24 h compared with vehicle control. Each dot represents a Reactome pathway associated with cytoskeletal organization or cellular mechanics. Dot colour represents the false discovery rate (FDR), and dot size corresponds to −log10(FDR). Positive NES values indicate enrichment in narciclasine-treated cells. The dashed vertical line marks NES = 0. **(B)** MCF10A cells were treated with vehicle (DMSO), 1 µM narciclasine, or pre-treated with 20 µM Y27632 (1 hour) followed by 1 µM narciclasine for 24 h. Entotic structures were visualized by immunofluorescence staining for β-catenin (membrane, green), LAMP1 (lysosomal compartments) and DAPI (nuclei, blue), followed by z-stack confocal microscopy. Representative 3D confocal images are shown. White arrows indicate entotic structures. Selected regions (boxed) are magnified, with corresponding orthogonal z-stack views displayed adjacent to each image. **(C)** Summary of quantification of entotic events in MCF10A cells treated from (B). Experiments were performed in biological triplicate (n = 3). Data are presented as mean ± SEM. Statistical analysis for (C) was performed using one-way ANOVA followed by Bonferroni post-hoc test.

**Supplementary Figure 10. Narciclasine increase PLS3 expression and promotes entotic structure formation.**

**(A)** Representative immunoblot images of PLS3 and tubulin (loading control) in MCF7 treated with vehicle or 1uM narciclasine and compared to non-treated (NT) conditions for 24 h and summary of densitometric quantification of PLS3 protein levels from immunoblot images. **(B)** Representative 3D confocal images of live MCF7 cells expressing PLS3-GFP and stained with SiR-Actin (red) and Hoechst (blue) for figure 8C. **(C)** Representative 3D confocal images of live MCF7 cells expressing GFP and stained with SiR-Actin (red) and Hoechst (blue). In B and C, white arrows indicate entotic structures. Boxed regions showing representative entotic structures are magnified, with corresponding orthogonal z-stack views and PC:DAPI images shown alongside each image. Shown are different Z positions. **(D)** Representative time-lapse images (phase contrast and Hoechst) for figure 8E showing induction of entosis in MCF7 cells expressing PLS3. For all confocal microscopy panels, orthogonal z-stack views and magnified insets are provided. Experiments were performed in biological triplicate (n = 3). Data for (A) are presented as mean ± SEM and are normalized to NT controls within each experiment (set to 100%). Statistical analysis was performed using a paired two-tailed Student’s t-test (**p < 0.01).

## REFERENCES

1. Overholtzer, M. & Brugge, J. S. The cell biology of cell-in-cell structures. Nat Rev Mol Cell Biol 9, 796–809 (2008).

2. Krishna, S. & Overholtzer, M. Mechanisms and consequences of entosis. Cell Mol Life Sci 73, 2379–2386 (2016).

3. Kim, S., Lee, D., Kim, S. E. & Overholtzer, M. Entosis: the core mechanism and crosstalk with other cell death programs. Exp Mol Med 56, 870–876 (2024).

4. Gaptulbarova, K. А., et al. Mechanisms and significance of entosis for tumour growth and progression. Cell Death Discov. 10, 109 (2024).

5. Wang, X. et al. Cell-in-Cell Phenomenon and Its Relationship With Tumor Microenvironment and Tumor Progression: A Review. Front. Cell Dev. Biol. 7, (2019).

6. Zhang, X. et al. Subtype-Based Prognostic Analysis of Cell-in-Cell Structures in Early Breast Cancer. Front Oncol 9, 895 (2019).

7. Schwegler, M. et al. Prognostic Value of Homotypic Cell Internalization by Nonprofessional Phagocytic Cancer Cells. Biomed Res Int 2015, 359392 (2015).

8. Mackay, H. L. et al. Genomic instability in mutant p53 cancer cells upon entotic engulfment. Nat Commun 9, 3070 (2018).

9. Hayashi, A. et al. Genetic and clinical correlates of entosis in pancreatic ductal adenocarcinoma. Mod Pathol 33, 1822–1831 (2020).

10. Dolma, L. et al. Mutant p53 induces SH3BGRL expression to promote cell engulfment. Cell Death Discov 11, 288 (2025).

11. Gutwillig, A. et al. Transient cell-in-cell formation underlies tumor relapse and resistance to immunotherapy. Elife 11, e80315 (2022).

12. Julian, L. & Olson, M. F. Rho-associated coiled-coil containing kinases (ROCK): Structure, regulation, and functions. Small GTPases 5, e29846 (2014).

13. Tang, Y. et al. LncRNAs regulate the cytoskeleton and related Rho/ROCK signaling in cancer metastasis. Mol Cancer 17, 77 (2018).

14. Durgan, J. & Florey, O. Cancer cell cannibalism: Multiple triggers emerge for entosis. Biochimica et Biophysica Acta (BBA) - Molecular Cell Research 1865, 831–841 (2018).

15. Hamann, J. C. et al. Entosis Is Induced by Glucose Starvation. Cell Rep 20, 201–210 (2017).

16. Overholtzer, M. et al. A Nonapoptotic Cell Death Process, Entosis, that Occurs by Cell-in-Cell Invasion. Cell 131, 966–979 (2007).

17. Durgan, J. et al. Mitosis can drive cell cannibalism through entosis. eLife 6, e27134.

18. Chen, R., Ram, A., Albeck, J. G. & Overholtzer, M. Entosis is induced by ultraviolet radiation. iScience 24, 102902 (2021).

19. Bozkurt, E. et al. TRAIL signaling promotes entosis in colorectal cancer. J Cell Biol 220, e202010030 (2021).

20. Priya, R. et al. Feedback regulation through myosin II confers robustness on RhoA signalling at E-cadherin junctions. Nat Cell Biol 17, 1282–1293 (2015).

21. Hughes, C. S. et al. Single-pot, solid-phase-enhanced sample preparation for proteomics experiments. Nat Protoc 14, 68–85 (2019).

22. Moggridge, S., Sorensen, P. H., Morin, G. B. & Hughes, C. S. Extending the Compatibility of the SP3 Paramagnetic Bead Processing Approach for Proteomics. J Proteome Res 17, 1730–1740 (2018).

23. Hsiao, Y. et al. Analysis and Visualization of Quantitative Proteomics Data Using FragPipe-Analyst. J Proteome Res 23, 4303–4315 (2024).

24. Ritchie, M. E. et al. limma powers differential expression analyses for RNA-sequencing and microarray studies. Nucleic Acids Res 43, e47 (2015).

25. Yu, G., Wang, L.-G., Han, Y. & He, Q.-Y. clusterProfiler: an R Package for Comparing Biological Themes Among Gene Clusters. OMICS 16, 284–287 (2012).

26. Paul-Gilloteaux, P. et al. eC-CLEM: flexible multidimensional registration software for correlative microscopies. Nat Methods 14, 102–103 (2017).

27. Perez-Riverol, Y. et al. The PRIDE database at 20 years: 2025 update. Nucleic Acids Res 53, D543–D553 (2025).

28. Lefranc, F. et al. Narciclasine, a plant growth modulator, activates Rho and stress fibers in glioblastoma cells. Mol Cancer Ther 8, 1739–1750 (2009).

29. Garanina, A. S. et al. Consecutive entosis stages in human substrate-dependent cultured cells. Sci Rep 7, 12555 (2017).

30. Florey, O., Kim, S. E., Sandoval, C. P., Haynes, C. M. & Overholtzer, M. Autophagy machinery mediates macroendocytic processing and entotic cell death by targeting single membranes. Nat Cell Biol 13, 1335–1343 (2011).

31. Kovács, M., Tóth, J., Hetényi, C., Málnási-Csizmadia, A. & Sellers, J. R. Mechanism of Blebbistatin Inhibition of Myosin II *. Journal of Biological Chemistry 279, 35557–35563 (2004).

32. Sebbagh, M. et al. Caspase-3-mediated cleavage of ROCK I induces MLC phosphorylation and apoptotic membrane blebbing. Nat Cell Biol 3, 346–352 (2001).

33. Győrffy, B. Survival analysis across the entire transcriptome identifies biomarkers with the highest prognostic power in breast cancer. Comput Struct Biotechnol J 19, 4101–4109 (2021).

34. Sun, Q., Cibas, E. S., Huang, H., Hodgson, L. & Overholtzer, M. Induction of entosis by epithelial cadherin expression. Cell Res 24, 1288–1298 (2014).

35. Wang, M. et al. Mechanical Ring Interfaces between Adherens Junction and Contractile Actomyosin to Coordinate Entotic Cell-in-Cell Formation. Cell Rep 32, 108071 (2020).

36. Sun, Q. et al. Competition between human cells by entosis. Cell Res 24, 1299–1310 (2014).

37. Bräutigam, J. et al. Narciclasine inhibits angiogenic processes by activation of Rho kinase and by downregulation of the VEGF receptor 2. Journal of Molecular and Cellular Cardiology 135, 97–108 (2019).

38. Stuelten, C. H., Parent, C. A. & Montell, D. J. Cell motility in cancer invasion and metastasis: insights from simple model organisms. Nat Rev Cancer 18, 296–312 (2018).

39. Ko, E., Kim, D., Min, D. W., Kwon, S.-H. & Lee, J.-Y. Nrf2 regulates cell motility through RhoA–ROCK1 signalling in non-small-cell lung cancer cells. Sci Rep 11, 1247 (2021).

40. Lochhead, P. A., Wickman, G., Mezna, M. & Olson, M. F. Activating ROCK1 somatic mutations in human cancer. Oncogene 29, 2591–2598 (2010).

41. Shinomiya, H. Plastin Family of Actin-Bundling Proteins: Its Functions in Leukocytes, Neurons, Intestines, and Cancer. Int J Cell Biol 2012, 213492 (2012).

42. Yokobori, T. et al. Plastin3 is a novel marker for circulating tumor cells undergoing the epithelial-mesenchymal transition and is associated with colorectal cancer prognosis. Cancer Res 73, 2059–2069 (2013).

43. Park, S. Y. et al. T-plastin contributes to epithelial-mesenchymal transition in human lung cancer cells through FAK/AKT/Slug axis signaling pathway. BMB Rep 57, 305–310 (2024).

44. Ueo, H. et al. Circulating tumour cell-derived plastin3 is a novel marker for predicting long-term prognosis in patients with breast cancer. Br J Cancer 112, 1519–1526 (2015).

45. Morley, S. C. The Actin-Bundling Protein L-Plastin: A Critical Regulator of Immune Cell Function. Int J Cell Biol 2012, 935173 (2012).

46. Machado, R. A. C. et al. L-plastin Ser5 phosphorylation is modulated by the PI3K/SGK pathway and promotes breast cancer cell invasiveness. Cell Commun Signal 19, 22 (2021).

47. Li, J. & Zhao, R. Expression and clinical significance of L-plastin in colorectal carcinoma. J Gastrointest Surg 15, 1982–1988 (2011).

48. Seo, S. et al. A novel protein LZTFL1 regulates ciliary trafficking of the BBSome and Smoothened. PLoS Genet 7, e1002358 (2011).

49. Wei, Q. et al. Tumor Suppressive Functions of Leucine Zipper Transcription Factor Like 1. Cancer Res 70, 2942–2950 (2010).

50. Grewal, T. et al. Annexin A6—A multifunctional scaffold in cell motility. Cell Adh Migr 11, 288–304 (2017).

51. Demonbreun, A. R. et al. Recombinant annexin A6 promotes membrane repair and protects against muscle injury. J Clin Invest 129, 4657–4670 (2019).

52. Sakwe, A. M., Koumangoye, R., Guillory, B. & Ochieng, J. Annexin A6 contributes to the invasiveness of breast carcinoma cells by influencing the organization and localization of functional focal adhesions. Exp Cell Res 317, 823–837 (2011).

53. Wolff, L. et al. Plastin 3 in health and disease: a matter of balance. Cell Mol Life Sci 78, 5275–5301 (2021).

54. Mei, L. et al. Structural mechanism for bidirectional actin cross-linking by T-plastin. Proceedings of the National Academy of Sciences 119, e2205370119 (2022).

55. Garbett, D. et al. T-Plastin reinforces membrane protrusions to bridge matrix gaps during cell migration. Nat Commun 11, 4818 (2020).

56. Xue, F., Janzen, D. M. & Knecht, D. A. Contribution of Filopodia to Cell Migration: A Mechanical Link between Protrusion and Contraction. Int J Cell Biol 2010, 507821 (2010).

57. Lee, A. R. & Park, C. Y. Orai1 is an Entotic Ca2+ Channel for Non-Apoptotic Cell Death, Entosis in Cancer Development. Adv Sci (Weinh) 10, e2205913 (2023).

58. Hinojosa, L. S., Holst, M., Baarlink, C. & Grosse, R. MRTF transcription and Ezrin-dependent plasma membrane blebbing are required for entotic invasion. J Cell Biol 216, 3087–3095 (2017).

